# A dedicated hypothalamic oxytocin circuit controls aversive social learning

**DOI:** 10.1101/2022.12.14.519639

**Authors:** Takuya Osakada, Rongzhen Yan, Yiwen Jiang, Dongyu Wei, Rina Tabuchi, Bing Dai, Xiaohan Wang, Gavin Zhao, Clara Xi Wang, Richard W. Tsien, Adam C. Mar, Dayu Lin

## Abstract

To survive and thrive in a complex social group, it is essential to not only know who to approach but more importantly who to avoid. After a single defeat, mice learn to stay away from the winning aggressor for weeks. Here, we identify oxytocin neurons in the retrochiasmatic supraoptic nucleus (SOR^OXT^) and oxytocin receptor expressing cells in the anterior subdivision of ventromedial hypothalamus, ventrolateral part (aVMHvl^OXTR^) as a key circuit motif for defeat-induced social avoidance learning. After defeat, aVMHvl^OXTR^ cells drastically increase their responses to aggressor cues. This response change is functionally important as optogenetic activation of aVMHvl^OXTR^ cells elicits time-locked social avoidance towards a benign social target whereas inactivating the cells suppresses defeat-induced social avoidance. Furthermore, OXTR in the aVMHvl is itself essential for the behavior change. Knocking out OXTR in the aVMHvl or antagonizing the receptor during defeat, but not during post-defeat social interaction, impairs defeat-induced social avoidance. aVMHvl^OXTR^ receives its private source of oxytocin from SOR^OXT^ cells, which are highly activated by the noxious somatosensory inputs associated with defeat. Oxytocin released from SOR^OXT^ depolarizes aVMHvl^OXTR^ cells and facilitates their synaptic potentiation, and hence, increases aVMHvl^OXTR^ cell responses to aggressor cues. Ablating SOR^OXT^ cells impairs defeat-induced social avoidance learning whereas activating the cells promotes social avoidance after a subthreshold defeat experience. Altogether, our study reveals an essential role of SOR^OXT^-aVMHvl^OXTR^ circuit in defeat-induced social learning and highlights the importance of brain oxytocin system in social plasticity.

## Introduction

In the wild, resources such as food, territory, and potential mates are limited. Consequently, animals often need to compete for obtaining such resources, and fighting is a major means to achieve that^1^. As a fight progresses, the animal with superior physical strength and fighting ability starts to dominate, initiating most of the offensive attacks and actively pursuing its opponent. Meanwhile, the losing animal starts to act more defensively, focusing on escaping from the opponent’s attacks and minimizing physical damage^2, 3^. After fighting is terminated, typically by retreat of the loser, the traumatic defeat experience is clearly remembered. The loser continuously avoids close interaction with the winner and readily flight away when confronted^4^. In male mice, a single 10-minute defeat can induce avoidance towards the winner for up to 15 days^5^. Additionally, the loser learns to stay away from the area where defeat had occurred to presumably minimize the chance of encountering the aggressor^6, 7^. The defeat-induced avoidance is observed across species, including humans^8^. In the US, a quarter of kids between 12-18 years old reportedly experienced bullying and showed increased social isolation and school avoidance^9^.

Despite being a widespread and robust phenomenon, the neural mechanisms underlying the fast and long-lasting behavior changes induced by social defeat remain incompletely understood. Early studies mainly focused on conditioned defeat, a phenomenon describing a complete loss of fighting ability after defeat in male hamsters^10^. These studies concluded that defeat and non-social aversive experience, e.g. foot shock, utilizes the same brain circuit, including prefrontal cortex, basolateral amygdala, and hippocampus, for associative fear learning^11–13^. More recently, several studies suggest a potential role of ventromedial hypothalamus, ventrolateral part (VMHvl) in social defense and avoidance^6, 14–17^. VMHvl is a part of social behavior network and exclusively activated by conspecific cues^18–20^. Sakurai et al. captured VMHvl cells that are activated during defeat and found that re-activation of those cells elicited fear responses towards a benign conspecific^14^. Conversely, broad inactivation of the VMHvl and its surrounding area reduces social avoidance towards the aggressor one day after defeat^6^. Our previous study further revealed functional heterogeneity within the VMHvl — anterior VMHvl (aVMHvl) is preferentially activated during defeat while the posterior VMHvl (pVMHvl) is most activated during attack^15^. Optogenetic activation of the aVMHvl cells elicits freezing, upright postures, and avoidance of a conspecific, while activating the pVMHvl cells elicits approach, close investigation, and attack of a social target^15, 19^. These studies support a role of the aVMHvl in social avoidance and fear. However, whether the aVMHvl mediates defeat-induced behavior change towards the winner and the underlying neural process remains unknown. Here, we investigated this question using a series of recording, functional and molecular tools and found that aVMHvl cells that express oxytocin receptor (aVMHvl^OXTR^) undergo dramatic changes during defeat with the help of a private source of oxytocin to mediate defeat-induced social avoidance to the bully.

## Results

### One day social defeat induces social avoidance and fear

We used the social interaction (SI) test^21^ to characterize the behavior changes induced by defeat. During SI test, a C57BL/6 (C57) male or female mouse freely interacted with a cupped aggressive Swiss Webster (SW) male or lactating female mouse^22^ for 10 min (**Extended Data Fig. 1a-b**). The SI tests were done one-day before and one-day after a 10-min resident-intruder (RI) test during which the C57 test mouse was introduced into the home cage (HC) of the aggressor, the same as the one in SI tests, for 10 min (**Extended Data Fig. 1c**). During the RI test, SW aggressors attacked and defeated the C57 intruders quickly and repeatedly (**Extended Data Fig. 1d-e**). After several bouts of defeat, C57 intruder mice spent more time immobile and often stayed in corners (**Extended Data Fig. 1f-h**). During pre-defeat SI test, C57 test mice repeatedly approached and investigated the cupped aggressor and spent approximately half of time around the cup (**Extended Data Fig. 1i-p, 1s-w**). After defeat, the animal spent most time staying at the end far from the aggressor and less time investigating the aggressor (**Extended Data Fig. 1l-n and s-u**). The reduced total investigation time is due to a combined effect of reduced approach frequency and shorter investigation time per visit (**Extended Data Fig. 1o-p and v-w**). Additionally, when the C57 was at the far end from the aggressor, it significantly reduced its movement velocity, i.e. freezing more (**Extended Data Fig. 1q-r and x-y**). These results suggest that a single 10-min defeat is sufficient to induce both social avoidance—measured by the reduced interaction time with the SW, and social fear—measured by the increased immobility when the aggressor is far away. The defeat-induced behavior change was qualitatively similar between males and females although the extent of avoidance was lower in females than in males (**Extended Data Fig. 1l-p and s-w**).

To understand whether the defeat-induced avoidance is specific to the aggressor, we compared the behaviors of the test C57 towards the SW aggressors, Balb/C (BC) non-aggressors and unfamiliar C57 mice using a multi-animal social interaction (MSI) test (**Extended Data Fig. 2a-b**). One day after defeat, the C57 test males spent significantly less time investigating or around the cupped SW aggressor and moved at a higher velocity when near the cupped SW aggressor (**Extended Data Fig. 2c, e-g**). In contrast, the interaction between the defeated C57 and an unfamiliar C57 or a previously encountered BC remained unchanged (**Extended Data Fig. 2c, e-g**). We also observed qualitatively similar results in C57 females mice during the MSI test (**Extended Data Fig. 2d, h-j**). Thus, one-time 10-min defeat induced winner-specific social avoidance.

### aVMHvl^OXTR^ cells in defeated animals increase responses to the winner of the fight

Our previous study showed that aVMHvl expressed a high level of c-Fos after defeat in comparison to that after winning^15^. Nasanbuyan et. al. reported that defeat-induced c-Fos overlaps well with OXTR at the VMHvl^23^. We thus examined defeat-induced c-Fos in OXTR^Cre^;Ai6 (OXTR^ZsGreen^) mice^24^ and found a significantly higher number of OXTR and c-Fos double positive cells (OXTR+Fos+) in the aVMHvl (Bregma: −1.36 mm to −1.52 mm) after defeat than that after attack while these two behaviors induced a similar number of OXTR+Fos+ cells in the pVMHvl (Bregma: −1.68 mm to −1.84 mm) (**Extended Data Fig. 3**). Consistent with the c-Fos expression pattern, fiber photometry recording of GCaMP6f expressing aVMHvl^OXTR^ cells in male mice (OXTR^GCaMP6^) revealed little activity increase of aVMHvl^OXTR^ cells during investigating or attacking a non-aggressive BC male intruder despite moderate initial response during intruder introduction (**Fig. 1a-c, f and i**). In contrast, when the test mouse fought and was defeated by an aggressive SW mouse, in either the test mouse’s or the SW aggressor’s cage, aVMHvl^OXTR^ cells strongly increased activity (**Fig. 1d-e, g-i**). Thus, aVMHvl^OXTR^ cells are activated during defeat but not winning regardless of the environment. Similar to the results in males, female aVMHvl^OXTR^ cells were also highly activated during defeat by a lactating SW female while showing no activity change during non-agonistic interactions with naïve C57 females (**Extended Data Fig. 4a-h**).

**Fig. 1:**
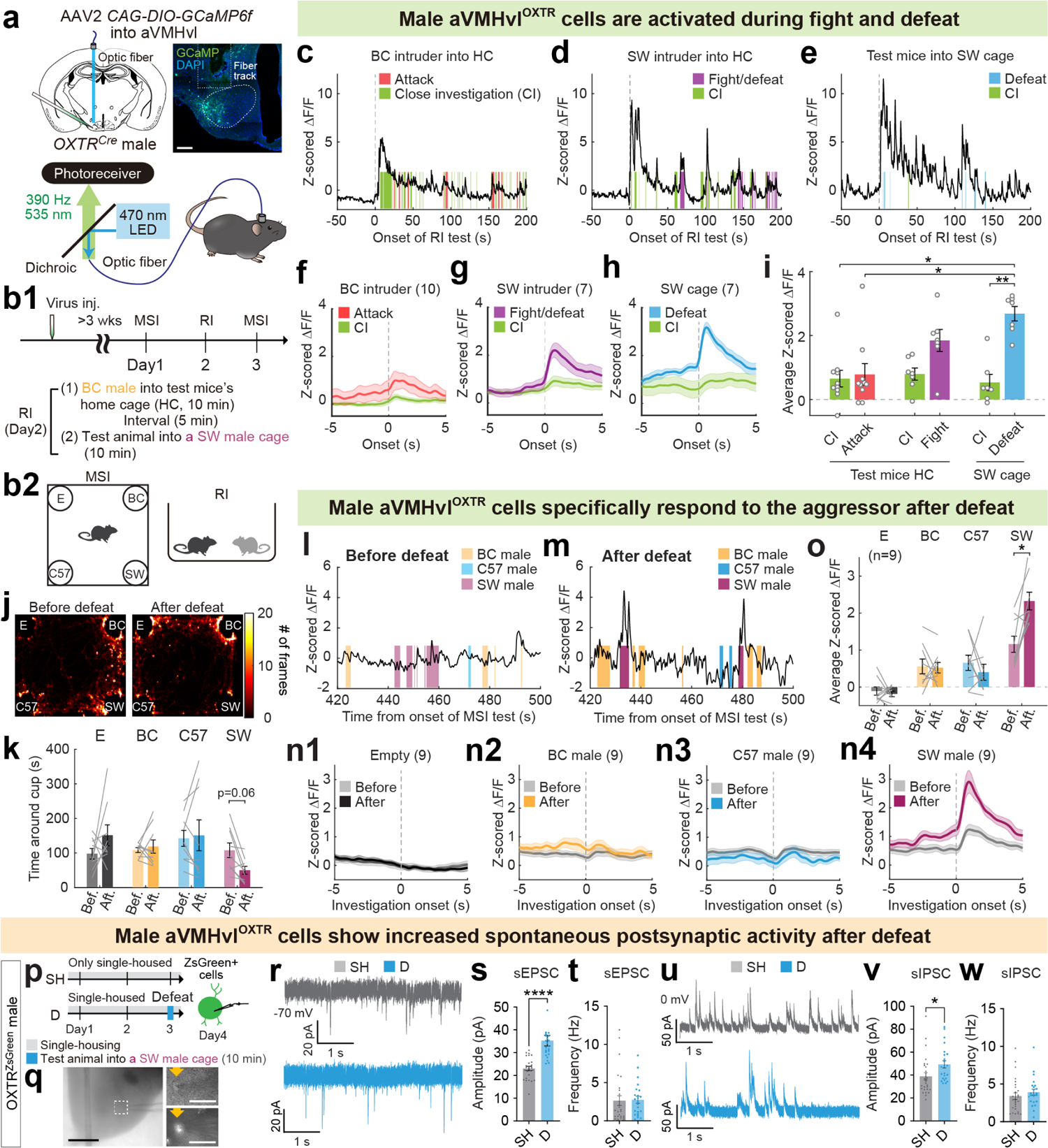
Male aVMHvl^OXTR^ cells significantly increase responses to aggressors after defeat. **a,** Schematics of virus injection and a representative histological image for the fiber photometry recording. Dashed line marks the aVMH. Scale bar: 200 µm. **b,** Experimental timeline. RI: resident-intruder test. MSI: multi-animal social interaction test. HC: home cage. **c-e,** Representative Z scored GCaMP6f traces of aVMHvl^OXTR^ cells from an animal that closely investigated (CI) and attacked a BC male intruder (**c**), an animal that fought and was later defeated by a SW aggressive male intruder (**d**), and an animal that was defeated by a resident SW male aggressor in the SW cage (**e**). **f-h**, Post-event histograms (PETHs) of Z scored GCaMP6f signals aligned to close investigation (CI) and attack of BC intruders, n=10 (**f**), CI and fighting/being defeated by SW intruders, n=7 (**g**), and CI and being defeated by aggressive SW residents, n=7 (**h**). **i,** Averaged Z scored ΔF/F responses of aVMHvl^OXTR^ cells during various social behaviors. **j,** Heatmaps showing the body center location of a recording mouse in MSI tests before and after defeat. **k,** Time the test animals spent around each cup during MSI tests before and after defeat. n=9. **l-m,** Representative Z scored GCaMP6f traces from an animal during pre-defeat (**l**) and post-defeat (**m**) MSI tests. Shades represent investigation periods of different cupped stimulus animals. Yellow: familiar BC male; Blue: unfamiliar C57 male; Maroon: familiar SW aggressor. Periods investigating the empty cup were not marked. **n1-n4,** PETHs of Z scored GCaMP6f signals aligned to onset of investigation of different cupped stimulus animals. Gray: pre-defeat; Color: post-defeat. **o,** Average Z scored ΔF/F of aVMHvl^OXTR^ cells during investigation of various cupped stimuli in the pre-defeat and post-defeat MSI tests. n=9. **p,** Experimental timeline of slice recording of aVMHvl^OXTR^ cells from single housed (SH) and defeated (D) male mice. **q**, Representative images show a recorded OXTR^ZsGreen^ cell within the aVMHvl. Right shows Infrared differential interference contrast (IR-DIC) (top) and fluorescent (bottom) images of the boxed area on the left. Yellow arrows indicate the recorded cell. Scale bars: 500 µm (left) and 50 µm (right). **r**, Representative sEPSC traces from cells in SH (top) and D (bottom) groups. Scale bars: 20 pA and 1 sec. **s-t**, Amplitude (**s**) and frequency (**t**) of sEPSCs of aVMHvl^OXTR^ cells recorded from SH and D groups. n= 25 cells from 3 mice for SH, and n= 23 cells from 3 mice for D group. **u**, Representative sIPSC traces from cells in SH (top) and D (bottom) groups. Scale bars: 50 pA and 1 sec. **v-w**, Amplitude (**v**) and frequency (**w**) of sIPSCs recorded from cells in SH and D groups. n= 27 cells from 3 mice for SH, and n= 23 cells from 3 mice for D group. Shades in **f-h, n** and error bars in **i, k, o, s, t, v,** and **w** represent ± SEM. Circles and lines in **i, k, o, s, t, v, and w** represent individual animals or recorded cells. Statistical analyses were performed with Kruskal-Wallis test with Dunn’s multiple comparisons test (**i**), two-way repeated-measure ANOVA with Sidak’s multiple comparisons test (**k, o**), Mann-Whitney test (**s, t**), and unpaired t-test (**v, w**). All statistical analyses were two-tailed. *p<0.05, **p<0.01, and ****p<0.0001.

After defeat, OXTR^GCaMP6^ mice showed less interaction with the cupped aggressor, supporting the emergence of social avoidance (**Fig. 1j-k, Extended Data Fig. 5a-g**). Strikingly, aVMHvl^OXTR^ cells showed a drastic increase in response to the SW aggressor after defeat (**Fig. 1l-o, Extended Data Fig. 5h-k**). During the pre-defeat SI test, little Ca^2+^ response was observed when the C57 OXTR^GCaMP6^ male mice investigated the cupped SW aggressor whereas aVMHvl^OXTR^ cells increased activity strongly during SW investigation post-defeat (**Extended Data Fig. 5h-k**). This defeat-induced response increase was specific to the winner of the fight as we observed no change in response to the previously encountered BC or an unfamiliar C57 male in the post-defeat MSI test (**Fig. 1l-o**). In females, aVMHvl^OXTR^ cells similarly increased response to the aggressive lactating SW females after defeat (**Extended Data Fig. 4m-p).** Our *in vivo* recording result is also consistent with the c-Fos expression pattern: interaction with the cupped aggressor for 10-minutes induced significantly higher level of c-Fos in the aVMHvl^OXTR^ cells, but not pVMHvl^OXTR^ cells, in animals that were defeated by the aggressor for 2 days in comparison to those that interacted with the cupped aggressor for 2 days (**Extended Data Fig. 6**).

To understand the physiological and synaptic changes that may underlie the *in vivo* response change, We performed patch-clamp recording of aVMHvl^OXTR^ cells in brain slices from OXTR^Cre^;Ai6 male mice that experienced a 10-min defeat one day prior to the recording **(Fig. 1p-q)**. Control cells were recorded from single-housed males that encountered no intruder **(Fig. 1p)**. We found that the magnitude of spontaneous excitatory postsynaptic current (sEPSC) of aVMHvl^OXTR^ cells in the defeated males (D) was significantly higher than that in single-housed males (SH) although the sEPSC frequency did not differ between these two groups (**Fig. 1r-t**). Spontaneous inhibitory postsynaptic currents (sIPSCs) also slightly increased in magnitude with no change in frequency (**Fig. 1u-w**). In contrast, the current-frequency relation of aVMHvl^OXTR^ cells, resting membrane potential, rheobase and membrane resistance did not differ between defeated and single housed males, suggesting no change in cell excitability **(Extended Data Fig. 7)**. These data suggest that synaptic potentiation occurred in aVMHvl^OXTR^ neurons after one-day defeat and likely contribute toward significantly increased *in vivo* responses to the aggressor.

### aVMHvl^OXTR^ cells are necessary and sufficient for defeat-induced social avoidance

What is the increased aVMHvl^OXTR^ cell response after defeat good for? To address this question, we expressed Channelrhodopsin-2 (ChR2)^25^ in the aVMHvl^OXTR^ cells (OXTR^ChR2^) and optogenetically activated the cells in undefeated animals during SI tests (**Fig. 2a-b**). Control animals expressed GFP in the aVMHvl^OXTR^ cells (OXTR^GFP^). Upon light delivery, OXTR^ChR2^, but not OXTR^GFP^ animals, showed robust avoidance of the cupped aggressor as indicated by significantly increased distance from the aggressor, reduced time spent investigating the aggressor, and decreased approach frequency (**Fig. 2c-f**). Activation of aVMHvl^OXTR^ cells also elicited strong social fear as indicated by a significantly increased immobility (**Fig. 2g**). The stimulation-induced avoidance and fear is not aggressive mouse-specific as similar behavior changes were observed when the cupped animal was a non-aggressive BC male (**Extended Data Fig. 8**). These results suggest that the increased aVMHvl^OXTR^ response to the winner after defeat is functionally relevant as high activity of the cells drives social avoidance and fear.

**Fig. 2:**
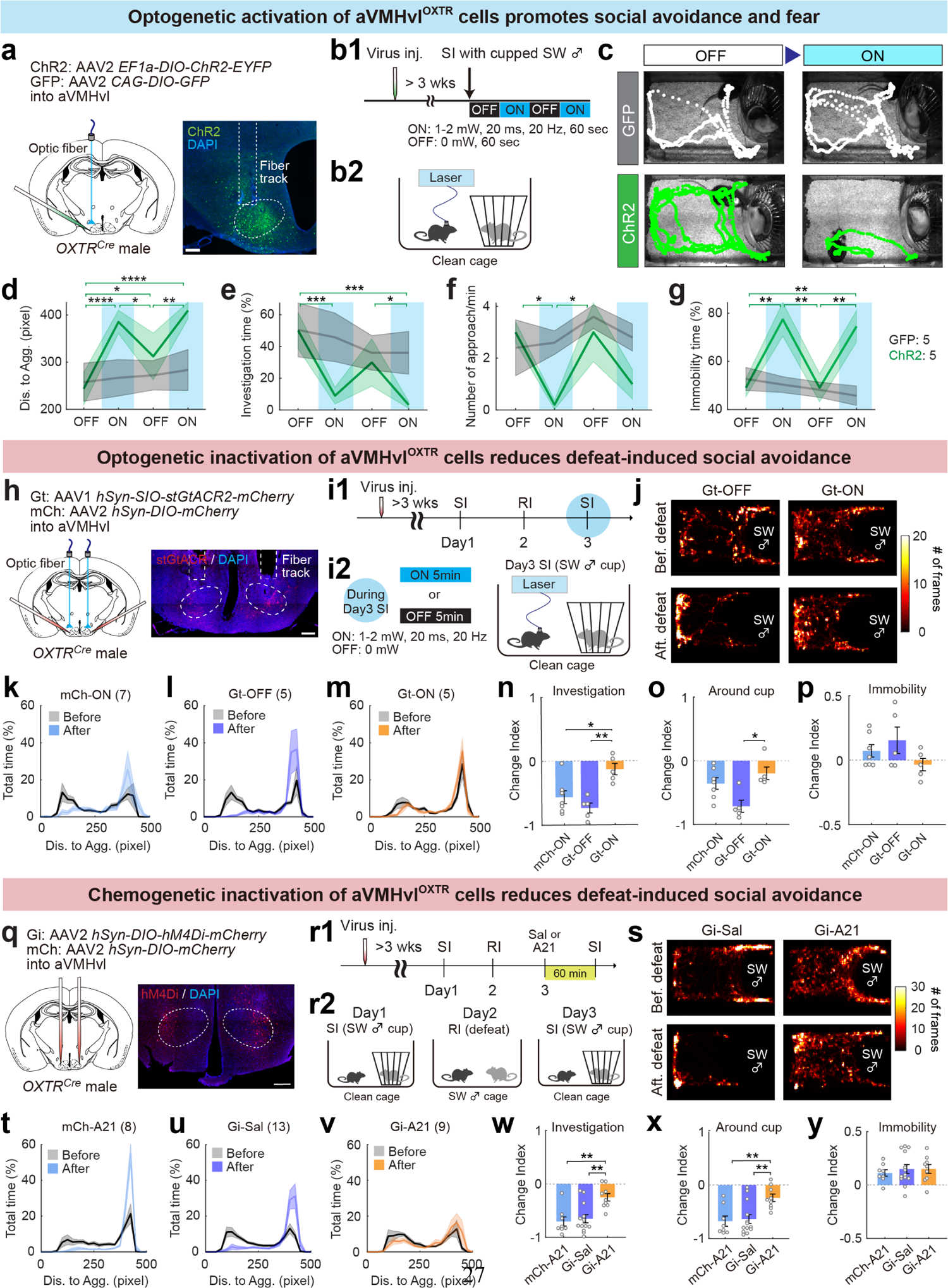
aVMHvl^OXTR^ cells are necessary and sufficient for post-defeat social avoidance. **a,** Schematics of virus injection and a representative histological image for the optogenetic activation experiment. Dashed line marks the aVMH. Scale bar: 200 µm. **b,** Experimental timeline and light delivery protocol (**b1**) and cartoon illustration of the behavior assay (**b2**). **c,** Video frames overlaid with representative movement trajectories of a GFP (top) and a ChR2 animal (bottom) during 60-s light-off (OFF) and light-on (ON) periods. **d-g,** Average distance to the aggressor cup (**d**), percentage of time spent investigating the cupped aggressor (**e**), frequency of approach (**f**), and the percentage of time spent immobile(**g**) during ON (blue shades) and OFF (no shade) periods in GFP (gray) and ChR2 groups. n=5 male mice in each group. Statistical results are between ON and OFF periods of ChR2 group. No significant difference in any parameters between ON and OFF periods in GFP groups. **h,** Schematics of virus injection and a representative histological image for the optogenetic inactivation experiment. Dashed lines mark bilateral aVMH. Scale bar: 200 µm. **i.** Experimental timeline (**i1**) and light delivery protocol (**i2**). **j.** Heatmaps showing the body center location of two Gt animals during SI tests before and after defeat. One animal received light and the other did not during the post-defeat SI test. **k-m,** Distribution of the distance between the test animal’s body center and cupped aggressor during the pre-defeat (black) and post-defeat SI tests for various groups. n=7 (mCh-ON), 5 (Gt-OFF) and 5 (Gt-ON) male mice. **n-p,** Change index of the percentage of time spent investigating the cupped aggressor (**n**), the percentage of time spent around (distance <250 pixels) the aggressor cup (**o**), and the percentage of time spent immobile when far (distance >300 pixels) from the aggressor cup (**p**) during post-defeat SI tests for various groups. Change index was calculated as (P_after_ – P_before_)/ (P_after_ + P_before_). P_before_ and P_after_ are values of the behavior parameter during SI test before and after defeat, respectively. n=7 (mCh-ON), 5 (Gt-OFF) and 5 (Gt-ON) male mice. **q,** Schematics of virus injection and a representative histological image for the chemogenetic inactivation experiment. Dashed lines mark bilateral aVMH. Scale bar: 200 µm. **r,** Experimental timeline (**r1**) and cartoon illustration of the SI-RI (defeat)-SI assay (**r2**). **s,** Heatmaps showing the body center location of two Gi animals during SI tests before (top) and after defeat (bottom). One animal was injected with saline (left) and the other (right) was injected with Agonist 21 (A21) before the post-defeat SI test. **t-v,** Distribution of the distance between the test animal’s body center and cupped aggressor during the pre-defeat (gray) and post-defeat SI tests for various groups. n=8 (Gi-A21), 13 (Gi-Sal), and 9 (Gi-A21). **w-y,** Change index of the percentage of time spent investigating the cupped aggressor (**n**), the percentage of time spent around the aggressor cup (**o**), and the percentage of time spent immobile when far from the aggressor cup (**p**) during post-defeat SI tests for various groups. n=8 (Gi-A21), 13 (Gi-Sal), and 9 (Gi-A21) male mice. Shades in **d-g**, **k-m** and **t-v** and error bars in **n-p** and **w-y** represent ± SEM. Circles in **n-p** and **w-y** represent individual animals. Statistical analyses were performed with two-way repeated-measure ANOVA with Holm-Sidak’s multiple comparisons test (**d-g**), Kruskal-Wallis test with Dunn’s multiple comparisons test (**o, x**), and one-way ANOVA with Tukey’s multiple comparisons test (**n, p, w, y**). All statistical analyses were two-tailed. *p<0.05, **p<0.01, ***p<0.001 and ***p<0.0001.

To understand whether the increased activity of aVMHvl^OXTR^ cells after defeat is necessary for the behavior change, we inhibited male aVMHvl^OXTR^ cells optogenetically during the post-defeat SI test (**Fig. 2h-i**). Specifically, we expressed stGtACR2^26^ in aVMHvl^OXTR^ cells (**Fig. 2h**). Three weeks later, the test animal went through a pre-defeat SI test and then was defeated by a SW aggressor one day later during a 10-min RI test (**Fig. 2i**). On the next day, we delivered light to inhibit the cells during a 5-min post-defeat SI test (OXTR^Gt-ON^) (**Fig. 2i**). Control animals either expressed stGtACR2 in aVMHvl^OXTR^ and received no light (OXTR^Gt-OFF^) or expressed mCherry in aVMHvl^OXTR^ and received light during the SI test (OXTR^mCh-ON^) (**Fig. 2h-i**). In comparison to control animals, OXTR^Gt-ON^ males spent more time surrounding and investigating the cupped aggressor (**Fig. 2j-o**) although the time spent immobile when the test animal was far from the aggressor did not differ significantly across groups (**Fig. 2p**).

As a complementary strategy, we chemogenetically inhibited aVMHvl^OXTR^ cells during post-defeat SI test. Specifically, we virally expressed hM4Di in the aVMHvl of OXTR^Cre^ mice (OXTR^hM4Di^) (**Fig. 2q**). Three weeks later, the animals were defeated by a SW aggressor during a 10-min RI test, and on the next day, we i.p. injected saline or Agonist 21^27^, a ligand for hM4Di (OXTR^Gi-Sal^ and OXTR^Gi-A^^21^) into the test animal, and subjected the animal to the SI test 60 min later (**Fig. 2r**). An additional control group was injected with mCherry expressing virus and Agonist 21 (OXTR^mCh-A^^21^) and went through the same behavior paradigm (**Fig. 2q-r**). In comparison to OXTR^Gi-Sal^ and OXTR^mCh-A^^21^ groups, OXTR^Gi-A^^21^ animals spent more time surrounding and investigating the cupped aggressor during the post-defeat SI test (**Fig. 2s-x**). However, when OXTR^Gi-A^^21^ animals stayed far from the cupped aggressor, they also showed increased immobility as that of control animals, suggesting that the manipulation did not significantly reduce social fear **(Fig. 2y**).

### An essential role of aVMHvl OXTR signaling during defeat in social avoidance learning

Is the oxytocin-OXTR signaling at the aVMHvl important for the defeat induced social avoidance and fear? To address this question, we next knocked out OXTR in the aVMHvl by bilaterally injecting Cre-GFP virus into OXTR^flox/flox^ male mice^28^ (OXTR^aVMHvl-KO^) (**Fig. 3e**). Control animals were injected with GFP virus (OXTR^aVMHvl-GFP^) (**Fig. 3e**). To confirm the successful OXTR knockout, we injected Cre-GFP and GFP viruses each into one side of aVMHvl of OXTR^flox/flox^ male mice and performed *in vitro* patch clamp recording of GFP positive cells (**Fig. 3a**). While 7/16 (44%) of GFP positive aVMHvl cells in GFP expressing side were depolarized by TGOT, a highly specific OXTR agonist, no GFP positive cell was responsive to TGOT in Cre-GFP expressing side, suggesting that OXTR has been effectively knocked out by Cre-GFP (**Fig. 3b-d**).

**Fig. 3:**
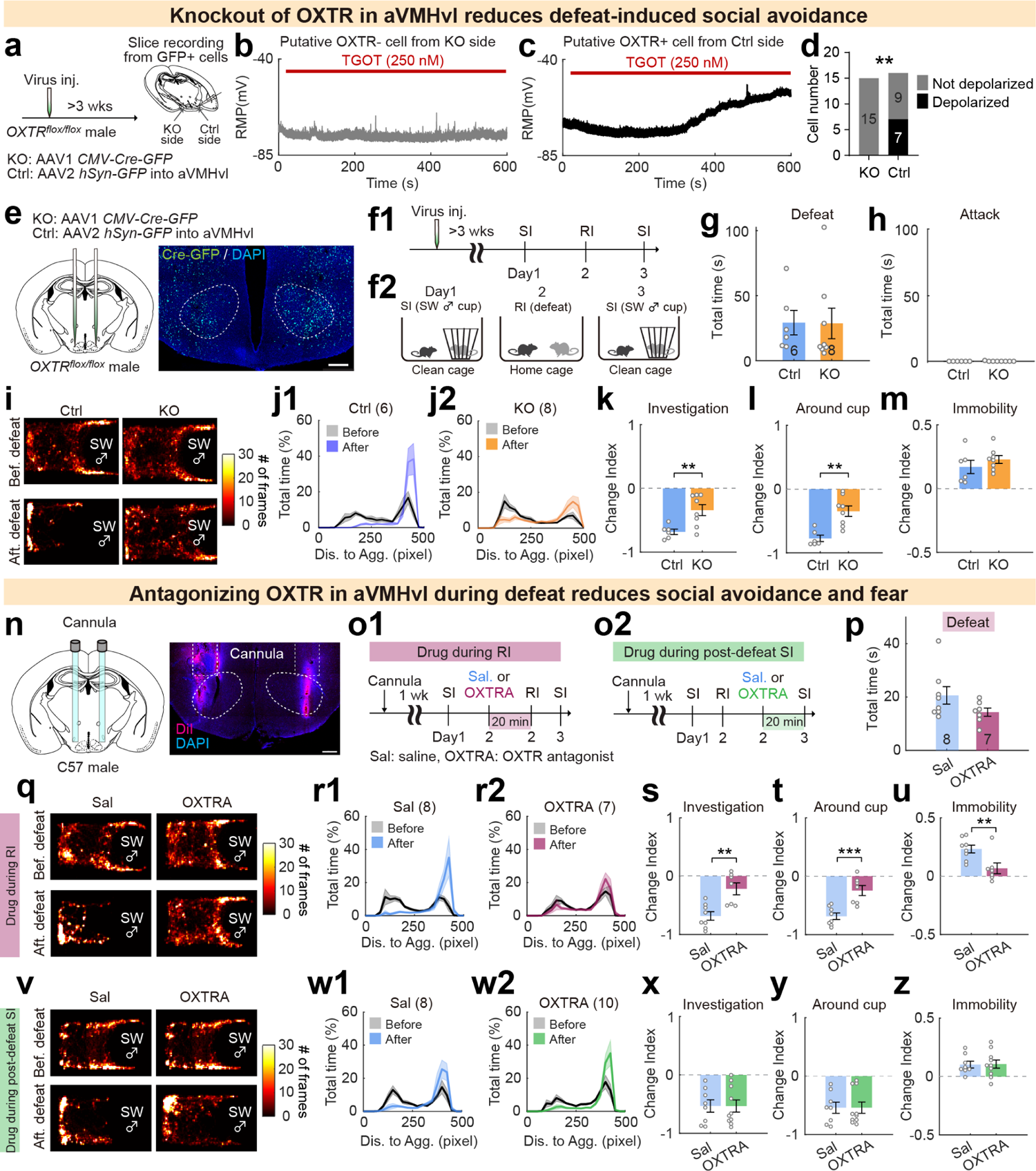
OXTRs in aVMHvl cells are essential for the acquisition of defeat-induced social avoidance. **a,** Experimental design to evaluate the efficiency of Cre-dependent knockout of OXTR in the aVMHvl. **b-c,** Representative recording traces of resting membrane potential (RMP) from GFP+ aVMHvl cells from knockout (KO) (**b**) and control (Ctrl) sides (**c**) under TGOT perfusion. Red bars represent periods of TGOT perfusion. **d,** The number of aVMHvl cells depolarized (>4mV) by TGOT in KO and control sides. n=15 cells (KO side) and 16 cells (control side) from 4 animals. **e,** Schematics of virus injection and a representative histological image showing Cre-GFP expression at the aVMHvl. Dashed lines mark bilateral aVMH. Scale bar: 200 µm. **f,** Experimental timeline (**f1**) and cartoon illustration of the SI-RI(defeat)-SI assay (**f2**). **g-h,** Total defeat (**g**) and attack (**h**) duration of the OXTR knockout and control mice in the 10-min RI tests with SW aggressors. n=6 (control) and 8 (KO) male mice. **i,** Heatmaps showing the body center location of an OXTR KO and a GFP control mouse during SI tests before and after defeat. **j,** Distribution of the distance between the test animal’s body center and cupped aggressor during the pre-defeat (gray) and post-defeat SI tests for GFP control (**j1**) and KO (**j2**) groups. n=6 (control) and 8 (KO) male mice. **k-m,** Change index of aggressor investigation time (**k**), time spent around the aggressor cup (**l**), and immobility (**m**) during post-defeat SI tests for various groups. n=6 (control) and 8 (KO) male mice. **n,** Schematics of cannulation and drug injection, and a representative histological image showing injection sites. Dashed lines mark bilateral aVMH. Scale bar: 200 µm. **o,** Experimental timelines for blocking OXTR with OXTR antagonist (OXTRA) during defeat (**o1**) and during post-defeat SI tests (**o2**). **p.** No significant difference in total defeat time of saline (Sal)-injected vs. OXTRA-injected animals during RI tests with SW aggressors. n=8 (Sal) and 7 (OXTRA) male mice. **q and v,** Heatmaps showing the body center location of representative mice during pre-defeat and post-defeat SI tests. The mice received saline or OXTRA injected 20 min prior to RI test (**q**) or prior to post-defeat SI test (**v**). **r and w,** Distribution of the distance between the test animal’s body center and cupped aggressor during the pre-defeat (gray) and post-defeat SI tests for animals that received saline (n=8, **r1**) or OXTRA (n=7, **r2**) injection prior to RI tests, and animals that received saline (n=8, **w1**) or OXTRA (n=10, **w2**) injection prior to SI tests. **s-u,** Change index of aggressor investigation time% (**s**), around the aggressor cup time% (**t**), and immobility time% (**u**) during post-defeat SI tests for animals that received saline (n=8) or OXTRA (n=7) injections prior to RI tests. **x-z,** graphs following the convention as **s-u**. OXTRA injection 20 min before post-defeat SI test did not change social avoidance in comparison to that after saline injection. Shades in **j, r and w** and error bars in **g, k-m, s-u, x-z** represent ± SEM. Circles in **g, h, k-m, s-u, and x-z** represent individual animals. Statistical analyses were performed with Fisher’s exact test (**d**), Mann Whitney test (**g, h, u, y**) and unpaired t-test (**k-m, p, s, t, x, z**). All statistical analyses were two-tailed. **p<0.01, ***p<0.001.

We then asked whether OXTR knockout at the aVMHvl affects defeat-induced social avoidance (**Fig. 3e-f**). During RI tests with SW aggressors, OXTR^aVMHvl-KO^ and OXTR^aVMHvl-GFP^ male mice were defeated for a similar amount of time and no test mouse in either group initiated attack towards the SW (**Fig. 3g-h**). In the post-defeat SI test, OXTR^aVMHvl-KO^ males spent more time surrounding and investigating the cupped aggressor than OXTR^aVMHvl-GFP^ males while both groups of animal showed increased immobility when far away from the cupped aggressor, supporting a main role of aVMHvl OXTR in defeat-induced social avoidance (**Fig. 3i-m**).

To understand whether the OXTR signaling is required for acquiring or expressing defeat-induced social avoidance, we injected L-368,899 hydrochloride (100 µM, 250 nL/side), a potent OXTR antagonist (OXTRA), into the aVMHvl of wildtype male mice either 20 min before the RI test (aVMHvl^OXTRA-RI^) or 20 min before the post-defeat SI test (aVMHvl^OXTRA-SI^) (**Fig. 3n-o**). Control males were injected with saline (aVMHvl^Sal-RI^ and aVMHvl^Sal-SI^) (**Fig. 3n-o**). OXTR antagonist injection prior to defeat did not affect aggressive behaviors of SW aggressors: aVMHvl^Sal-RI^ and aVMHvl^OXTRA-RI^ mice were defeated for a similar amount of time (**Fig. 3p**). During post-defeat SI test, aVMHvl^OXTRA-RI^ male mice spent significantly more time surrounding and investigating the cupped aggressor and reduced immobility when far away from the aggressor in comparison to aVMHvl^Sal-RI^ mice (**Fig. 3q-u**). In contrast, injecting OXTR antagonist before the post-defeat SI test had no effect on social avoidance or social fear: aVMHvl^OXTRA-SI^ and aVMHvl ^Sal-SI^ mice showed similarly high social avoidance and increased immobility (**Fig. 3v-z**). These results strongly suggest that OXTR signaling at the aVMHvl is necessary for social avoidance learning during defeat but not their expression during post-defeat social encounters.

### SOR is the main source of oxytocin for aVMHvl^OXTR^ cells

We next aimed to identify the source of oxytocin for aVMHvl^OXTR^ cells. Given that aVMHvl OXTR signaling is required during defeat for social avoidance learning, we reasoned that oxytocin release at the aVMHvl should occur during defeat. We examined the overlap between defeat-induced c-Fos and oxytocin and found that c-Fos and oxytocin double positive cells were present in paraventricular hypothalamic nucleus (PVN), supraoptic nucleus (SON) and retrochiasmatic supraoptic nucleus (SOR)^29^, a small region caudal to SON (and sometimes considered as a subdivision of SON^30, 31^) (**Fig. 4a-b**). SOR is particularly interesting as it contained the highest percentage (∼50%) of oxytocin cells that express defeat-induced c-Fos (**Fig. 4c**). Anatomically, SOR is located right next to the aVMHvl^OXTR^ cells (**Fig. 4d**), making it well positioned to provide OXT to aVMHvl cells through not only axonal release but also somatodendritic release^32, 33^. Vast majority of oxytocin cells express vesicular glutamate transporter 2 (Vglut2), suggesting that oxytocin cells may co-release glutamate (**Extended Data Fig. 9a-d**).

**Fig. 4:**
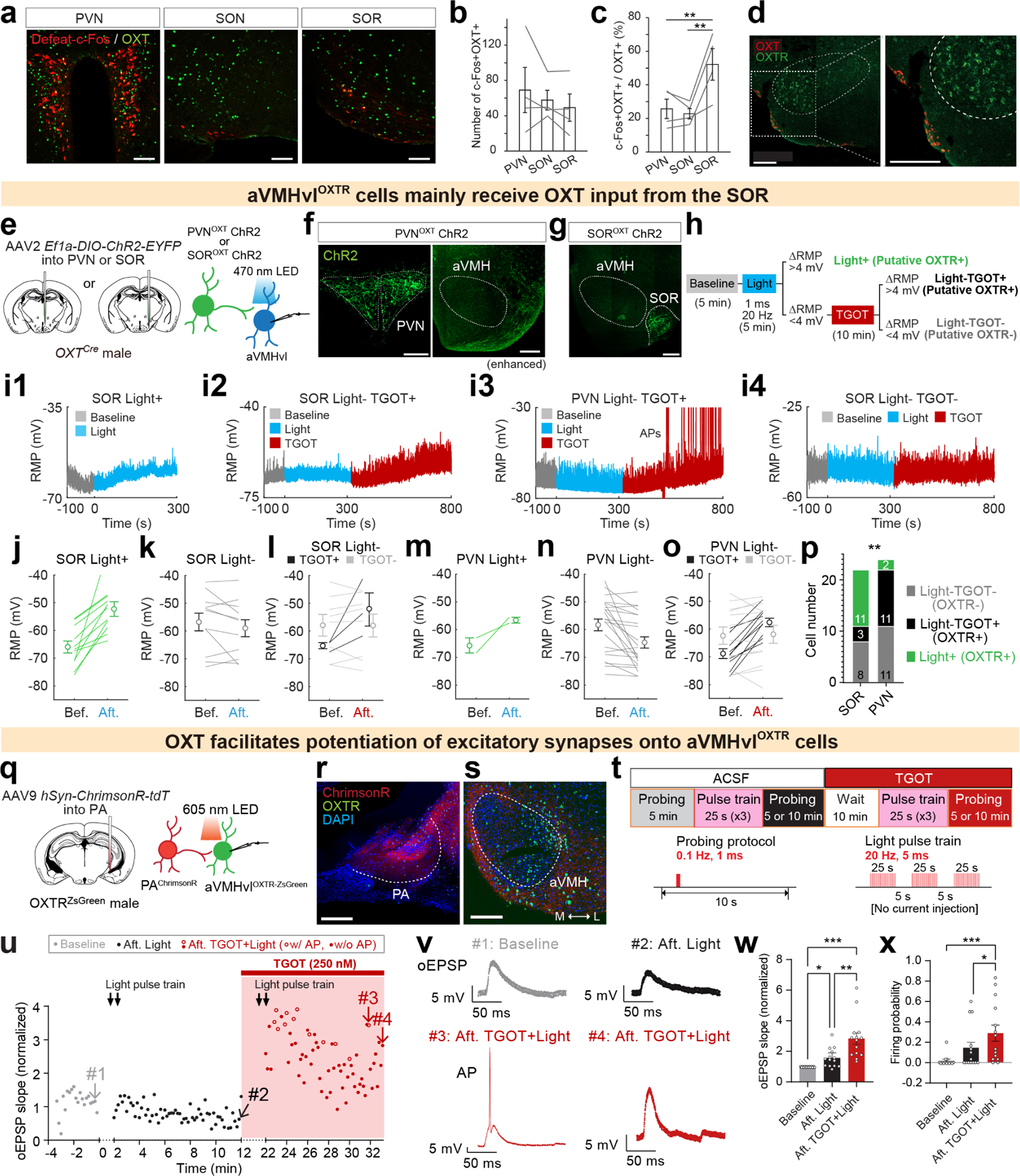
SOR is the main source of OXT for aVMHvl^OXTR^ cells. **a,** Representative immunohistochemical images showing expression of defeat-induced c-Fos (red) and oxytocin (OXT, green) in paraventricular hypothalamic nucleus (PVN), supraoptic nucleus (SON), and retrochiasmatic supraoptic nucleus (SOR) of male mice. Scale bar: 50 µm. **b,** Average number of c-Fos and OXT dual-positive cells in each region. Every other brain section was used for cell counting. (n=4 male mice) **c,** The percentage of c-Fos and OXT dual-positive cells in all OXT-positive cells in each region. (n=4 male mice) **d,** *In situ* hybridization images showing *OXT* (red) and *OXTR* (green) mRNA expression in the SOR and aVMHvl from a C57 male mouse. Dashed line marks the aVMH. Right image shows the enlarged view of the boxed area on the left. Scale bars: 200 µm. **e**, Experimental schematic showing slice recordings of aVMHvl cells while optogenetic activation of OXT inputs from the PVN or SOR. **f**, Representative images showing ChR2 expression in PVN^OXT^ cells and their axons surrounding the aVMH. The right image was digitally enhanced to reveal the florescence signal. Scale bars: 200 µm. **g**, ChR2 expression in SOR^OXT^ cells. Scale bar: 200 µm. **h,** Experimental design to examine aVMHvl cell responses to inputs from SOR^OXT^ or PVN^OXT^ cells and strategy to functionally determine OXTR expression of recorded cells. **i,** Representative recording traces showing RMPs at the baseline (gray), during light delivery (blue) and TGOT perfusion (red). **i1**, A putative OXTR+ cell that was depolarized by SOR^OXT^ light stimulation (SOR Light+); **i2**, a putative OXTR+ cell that was non-responsive to SOR^OXT^ activation but depolarized by TGOT (SOR Light-TGOT+); **i3**, a putative OXTR+ cell that was non-responsive to PVN^OXT^ terminal activation but depolarized by TGOT (PVN Light-TGOT+). Note that the cell started to fire action potentials (APs) after TGOT; **i4**, a putative OXTR-cell that did not respond to either SOR^OXT^ activation or TGOT perfusion (SOR Light-TGOT-). **j-k,** RMP change of aVMHvl cells that were depolarized (**j,** n=11 SOR Light+ cells) and not depolarized (**k,** n=11 SOR Light-cells) by SOR^OXT^ light activation. **l.** TGOT-induced RMP change of SOR Light-aVMHvl cells. 3 cells were depolarized after TGOT (SOR Light-TGOT+, black), while 8 were not (SOR Light-TGOT-, light gray). **m-n** RMP change of aVMHvl cells that were depolarized (**m,** PVN Light+, n=2 cells) and not depolarized (**n,** PVN Light-, n=22 cells) by PVN^OXT^ terminal activation. **o.** TGOT-induced RMP change of PVN Light-cells. 11 cells were depolarized after TGOT application (PVN Light-TGOT+, black) while 11 were not (PVN Light-TGOT-, light gray). **p,** Number of recorded cells in the aVMHvl that were Light-TGOT-(Putative OXTR-, gray), Light-TGOT+ (putative OXTR+, black), and Light+ (putative OXTR+, green). n=22 (SOR) and 24 (PVN) aVMHvl cells, each from 4 male mice. **q,** Schematics showing recording of aVMHvl^OXTR^ cells during optogenetic activation of posterior amygdala (PA) terminals. **r-s,** Representative images showing ChrimsonR in PA cells (**r**) and their axons around the aVMHvl (**s**). Scale bars: 200 µm. M: medial, L: lateral. **t,** Slice recording timeline (top), and light delivery protocols (bottom) for probing and inducing synaptic potentiation. **u,** Slopes of light evoked EPSP (oEPSP) of aVMHvl^OXTR^ cells at the baseline (gray), after light stimulation (black), after TGOT perfusion and light stimulation (red). Red bar indicates the period of TGOT perfusion. Open circles represent oEPSPs with action potential (AP) and closed circles represent those without AP. Gray arrows (#1-4) indicate representative oEPSPs shown in **v**. **v,** Representative oEPSPs during different recording phases. Scale bars, 5 mV and 50 ms. **w-x,** Normalized oEPSP slope (**w**) and light stimulation evoked firing probability (**x**) at baseline, after light stimulation without TGOT, and after light stimulation in the presence of TGOT. n=14 cells from 5 male mice. Error bars in **b, c, j-o, w and x** represent ± SEM. Circles and lines in **b, c, j-o, u, w and x** represent individual animals or recording cells. Statistical analyses were performed with one-way RM ANOVA with Tukey’s multiple comparisons test (**b-c, w**), Chi-square test (**p**), and Friedman test with Dunn’s multiple comparisons test (**x**). No statistical analysis was done for **j-o** given that the cells were categorized based on their response patterns. All statistical analyses were two-tailed. *p<0.05, **p<0.01, and ***p<0.001.

To understand the influence of oxytocin input on aVMHvl cell activity, we virally expressed ChR2 in SOR^OXT^ or PVN^OXT^ cells using OXT^Cre^ male mice^34^ and performed current clamp recording of aVMHvl cells on brain slices while delivering light pulses (1 ms, 20 Hz) for 5 min to activate SOR^OXT^ or PVN^OXT^ input (**Fig. 4e-g**). Upon activation of SOR^OXT^ input, 11/22 aVMHvl cells showed >4 mV increase in resting membrane potential, consistent with reported effect of OXT on VMHvl cells^35^, and we considered those cells as putatively OXTR positive (**Fig. 4h, i, j-k, and p**). In comparison, only 2/24 aVMHvl cells were obviously depolarized by PVN^OXT^ terminal activation (**Fig. 4m-n, p**). For aVMHvl cells that were not depolarized by PVN^OXT^ or SOR^OXT^ activation, we applied TGOT to functionally determine the cell’s expression of OXTR (**Fig. 4h**). We found that 3/11 of SOR^OXT^ activation unresponsive cells were depolarized by TGOT while 11/22 of PVN^OXT^ stimulation unresponsive cells did so (**Fig. 4h, i2-i4, l, o and p**). Altogether, we estimated that SOR^OXT^ and PVN^OXT^ inputs influence 80% (11/14 cells) and 15% (2/13 cells) of aVMHvl^OXTR^ cells, respectively. Although oxytocin cells likely co-express glutamate, we did not observe optogenetically evoked excitatory post-synaptic current (oEPSC) with light delivery to either SOR^OXT^ or PVN^OXT^ input (**Extended Data Fig. 9e-h**). These results suggest that SOR^OXT^ cells are the main source of oxytocin for aVMHvl cells and they do not influence aVMHvl^OXTR^ cell activity through fast neurotransmitter release.

We next asked whether oxytocin-OXTR signaling at the aVMHvl could facilitate synaptic potentiation as observed in defeated animals (**Fig. 1s**). To control the excitatory input to aVMHvl^OXTR^ cells, we virally expressed ChrimsonR^36^ in posterior amygdala (PA) cells and performed current clamp recording of aVMHvl^OXTR^ cells using OXTR^Cre^;Ai6 male mice (**Fig. 4q-s**). PA is the main extrahypothalamic glutamatergic input to the VMHvl and evokes strong monosynaptic EPSC from VMHvl cells^37–39^. For each recorded aVMHvl^OXTR^ cell, we probed its postsynaptic responses to PA inputs with 1 ms, 0.1 Hz 605 nm light pulses for 5 min (**Fig. 4t**). Then, we delivered 20 Hz, 5 ms, 25 s light pulses for 3 times to mimic the strong input from PA that could occur naturally, e.g. during interaction with a male conspecific^37^(**Fig. 4t**). After the light pulse train, we found that the postsynaptic response of aVMHvl^OXTR^ cells to the PA input only increased slightly (**Fig. 4u-w**). Strikingly, when the light pulses were delivered after bath application of TGOT for 10 min, the increase in light evoked excitatory postsynaptic potential (oEPSP) was significantly larger (**Fig. 4u-w**). As a result, aVMHvl^OXTR^ cells were more likely to fire action potentials upon 1-ms PA terminal stimulation (**Fig. 4x**). Thus, oxytocin-OXTR signaling at the aVMHvl can potentiate excitatory synaptic transmission onto aVMHvl^OXTR^ cells likely through a postsynaptic voltage-dependent mechanism^40^.

### SOR^OXT^ cells are specifically activated during social defeat, likely due to defeat-associated pain

To understand the *in vivo* response pattern of SOR^OXT^ cells, we expressed GCaMP6f in SOR^OXT^ cells using OXT^Cre^ male mice and performed fiber photometry recording of Ca^2+^ activity during pre- and post-defeat MSI tests as well as RI test with a non-aggressive BC intruder and a SW aggressor (**Fig. 5a-b**). We found that SOR^OXT^ cells are highly activated during each episode of fight or defeat with the SW aggressor, regardless of whether the defeat occurred in SW or test animal’s home cage, but showed no response when the recording animal investigated an opponent or attacked a BC intruder (**Fig. 5c-i**). However, in contrast to the increased response of aVMHvl^OXTR^ cells towards the aggressor after defeat, SOR^OXT^ cells showed no such increase in response (**Fig. 5j-o**). Indeed, SOR^OXT^ cells were not activated during any social target investigation regardless of the animal’s defeat experience (**Fig. 5j-o**).

**Fig. 5:**
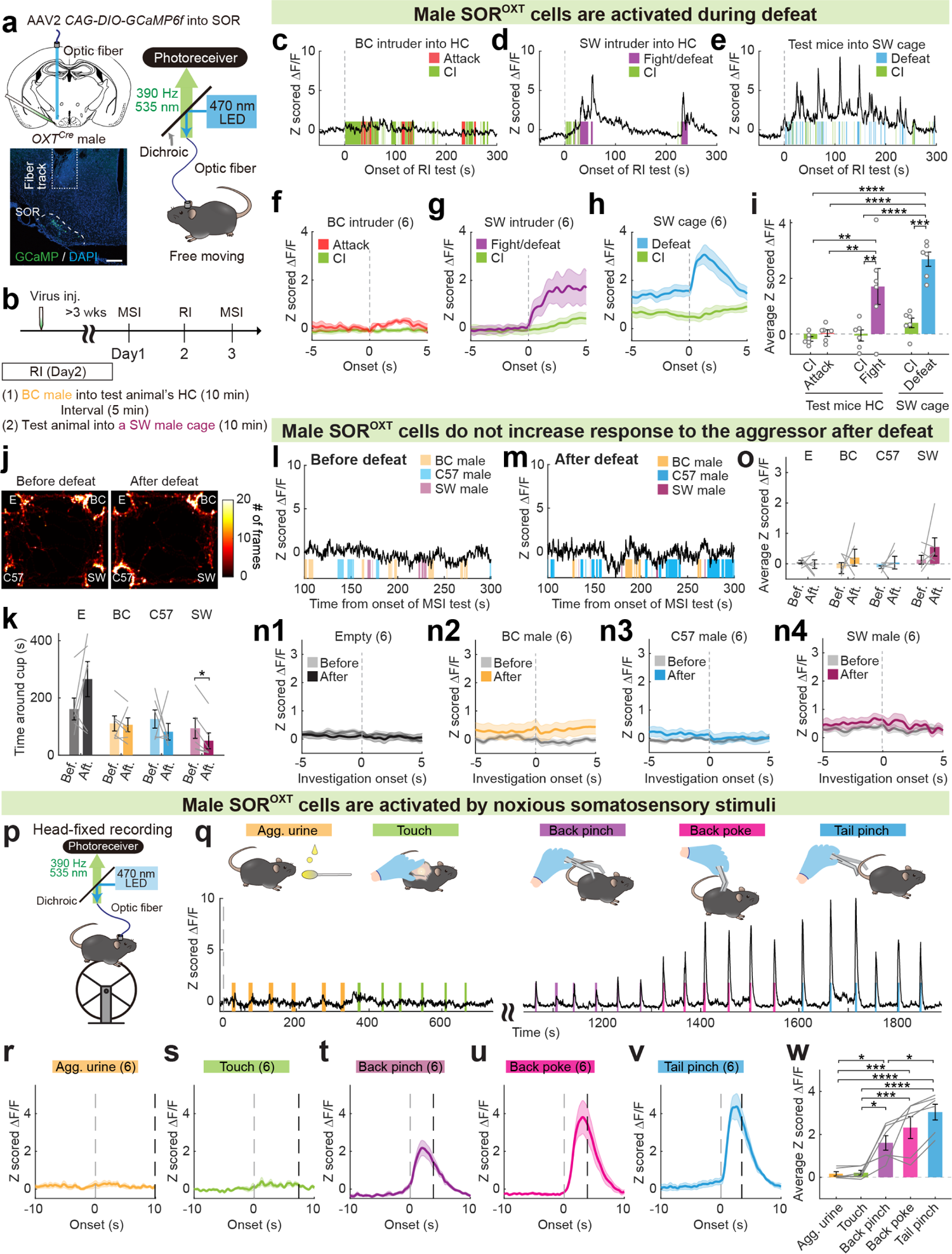
SOR^OXT^ cells are activated by noxious stimuli associated with defeat. **a,** Schematics of virus injection and a representative histological image for fiber photometry recording of SOR^OXT^ cells in males. Dashed line marks the SOR. Scale bar: 200 µm. **b,** Experimental timeline. **c-e,** Representative Z scored GCaMP6f traces of SOR^OXT^ cells from an animal that closely investigated (CI) and attacked a BC male intruder (**c**), an animal that fought and was later defeated by a SW aggressive male intruder (**d**), and an animal that was defeated by a resident SW male aggressor in the SW cage (**e**). **f-h**, PETHs of Z scored GCaMP6f signals aligned to CI and attack of BC intruders (**f**), CI and fighting/being defeated by SW intruders (**g**), and CI and being defeated by resident SW aggressors (**h**). n=6 male mice for each plot. **i,** Average Z scored ΔF/F responses of SOR^OXT^ cells during various social behaviors. **j,** Heatmaps showing the body center location of a recording mouse in MSI tests before and after defeat. E: empty cup, BC: BC non-aggressive male cup, C57: unfamiliar C57 cup, and SW: SW male aggressor cup. **k,** Time spent around each cup during MSI tests before and after defeat. Bef: before RI test; Aft: after RI test. n=6 male mice. **l-m,** Representative Z scored GCaMP6f traces from an animal during pre-defeat and post-defeat MSI tests. Shades represent investigation periods of different cupped stimulus animals. Periods investigating the empty cup were not marked. **n,** PETHs of Z scored GCaMP6f signals aligned to the onset of investigation of different cupped stimulus animals. Gray: pre-defeat; Color: post-defeat. n=6 male mice. **o,** Average Z scored ΔF/F responses of SOR^OXT^ cells during investigation of various cupped stimuli in the pre-defeat and post-defeat MSI tests. n=6 male mice. **p,** Schematics of head-fixed fiber photometry recording of SOR^OXT^ cells. **q,** Representative Z scored GCaMP6f trace of SOR^OXT^ cells during delivery of aggressor urine on a Q-tip (Urine), gentle touch, back pinch, back poke and tail pinch. **r-v,** PETHs of Z scored GCaMP6f signals aligned to onset of aggressor urine presentation (**r**), gentle touch (**s**), back pinch (**t**), back poke (**u**) and tail pinch (**v**). Gray and black dashes lines indicate the onset and average offset of stimulus presentation. **w,** Average Z scored ΔF/F during various stimulus presentation. Shades in **f-h, n** and **r-v**, and error bars in **i, k, o** and **w** represent ± SEM. Circles and lines in **i, k, o** and **w** represent individual animals. Statistical analyses were performed with one-way ANOVA with Tukey’s multiple comparisons test (**i**), two-way repeated measure ANOVA with Sidak’s multiple comparisons test (**k, o**), and one-way repeated measure ANOVA with Tukey’s multiple comparisons test (**w**). All statistical analyses were two-tailed. *p<0.05, **p<0.01, ***p<0.001, and ****p<0.0001.

Given the specific response of SOR^OXT^ cells during defeat, we hypothesized that SOR^OXT^ cells are activated by sensory inputs associated with being attacked, i.e. pain. To test this hypothesis, we recorded Ca^2+^ activity of SOR^OXT^ cells in head-fixed animals while presenting several sensory stimuli, including aggressor urine, gentle touch on the back, pinch and poke on the back, and tail pinch (**Fig. 5p-q**). SOR^OXT^ cells showed no response during gentle touch or urine presentation (**Fig. 5q-s**), moderate increase during back pinch (**Fig. 5t**), and robust and consistent activation during back poke and tail pinch (**Fig. 5u, v**). SOR^OXT^ cells in female mice showed similar response patterns as those in males (**Extended Data Fig. 10**). These results strongly suggest that among various social behaviors, SOR^OXT^ cells are activated specifically during defeat, likely due to the noxious somatosensory stimuli, e.g. pain, associated with the experience.

### SOR^OXT^ cells are functionally important for defeat-induced social avoidance

To understand the functional importance of SOR^OXT^ cells in defeat-induced social avoidance and fear learning, we virally expressed diphtheria toxin receptor (DTR) to ablate SOR^OXT^ cells using OXT^Cre^ male mice (OXT^DTR^) (**Fig. 6a**). The ablation was complete and specific as shown by a complete loss of oxytocin staining in the SOR in DTR-expressing but not GFP-expressing animals (OXT^GFP^) whereas vasopressin (AVP) expression in the SOR was not affected in OXT^DTR^ mice (**Fig. 6a-b**). During the RI test, OXT^DTR^ and OXT^GFP^ males were defeated for a similar amount of time by SW aggressors (**Fig. 6c-d**). However, during the post-defeat SI test, OXT^DTR^ mice spent more time around and investigating the cupped aggressor in comparison to OXT^GFP^ males (**Fig. 6e-i**) whereas OXT^DTR^ and OXT^GFP^ animals spent a similar amount time staying immobile when away from the aggressor (**Fig. 6j**). These results support a necessary role of SOR^OXT^ cells in defeat-induced social avoidance.

**Fig. 6:**
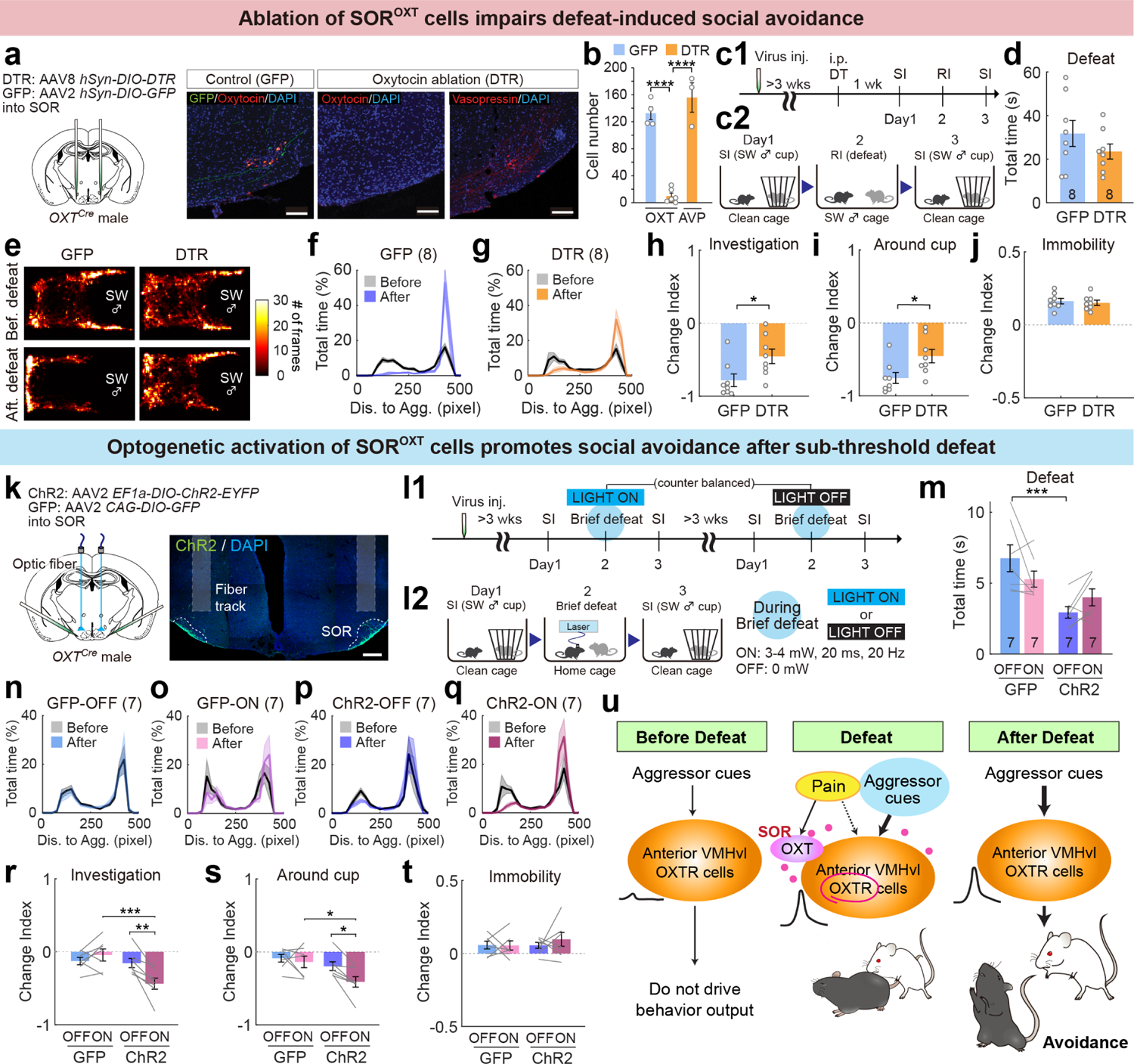
SOR^OXT^ cells are functionally essential for social avoidance learning. **a,** Schematics of virus injection and representative histological images showing OXT expression in the SOR of a GFP but not DTR mouse. Vasopressin (AVP) expression in the SOR remained in the DTR mouse. Scale bars: 100 µm. **b,** Number of OXT and AVP positive cells in the SOR in GFP (blue) and DTR (orange) mice. Every other brain section was used for cell counting. **c,** Experimental timeline (**c1**) and cartoon illustration of the SI-RI(defeat)-SI assay (**c2**). Diphtheria toxin (DT) was administrated into all the animals three weeks after virus injection. **d,** No significant difference in total defeat time experienced by GFP and DTR animals during the RI test with SW aggressors. n=8 male mice for each group. **e,** Heatmaps showing the body center location of representative GFP and DTR mice during pre-defeat and post-defeat SI tests. **f-g,** Distribution of the distance between the test animal’s body center and cupped aggressor during the pre-defeat (gray) and post-defeat SI tests for GFP (**f,** n=8) and DTR (**g,** n=8) animals. **h-j,** Change index of aggressor investigation time% (**h**), around the aggressor cup time% (**i**), and immobility time% (**j**) during post-defeat SI tests for GFP (n=8) and DTR (n=8) animals. **k,** Schematics of virus injection and a representative histology image for optogenetic activation of SOR^OXT^ cells. Scale bar: 200 µm. **l,** Experimental timeline (**l1**) and light delivery protocol during the RI test when animal experienced brief defeat (∼5s) (**l2**). **m,** Total defeat time of various groups during the RI test with SW aggressors. n=7 male mice for each group. **n-q,** Distribution of the distance between the test animal’s body center and cupped aggressor during the pre-defeat (gray) and post-defeat SI tests for GFP-light OFF (**n**), GFP-light ON (**o**), ChR2-light OFF (**p**), and ChR2-light ON groups (**q**). n= 7 male mice in each group. **r-t,** Change index of aggressor investigation time% (**r**), around the aggressor cup time% (**s**), and immobility time% (**t**) during post-defeat SI tests for various groups. **u.** A model for SOR^OXT^-aVMHvl^OXTR^ oxytocin signaling-dependent social avoidance learning after defeat. Before defeat, aVMHvl^OXTR^ cells do not respond to aggressor and no avoidance. During defeat, biting-induced pain evokes strong activation of SOR^OXT^ cells and release of oxytocin, which depolarizes aVMHvl^OXTR^ cells and potentiates the synaptic connections that carry the aggressor information. After defeat, when the animal encounters the aggressor again, due to the strengthened synaptic inputs, aVMHvl^OXTR^ cells are now activated and drive social avoidance. Shades in **f, g** and **n-q**, and error bars in **b, d, h-j, m,** and **r-t** represent ± SEM. Circles and lines in **b, d, h-j, m, and r-t** represent individual animals. Statistical analyses were performed with one-way ANOVA with Tukey’s multiple comparisons test (**b**), unpaired t-test (**d, i, j**), Mann-Whitney test (**h**), and two-way repeated measure ANOVA with Sidak’s multiple comparisons test (**m, r-t**). All statistical analyses were two-tailed. *p<0.05, **p<0.01, ***p<0.001 and ****p<0.0001.

Lastly, we asked whether activation of SOR^OXT^ cells, which naturally happens during defeat, could facilitate social avoidance. We virally expressed ChR2 or GFP in SOR^OXT^ cells using OXT^Cre^ male mice (OXT^ChR2^ and OXT^GFP^) and subjected the animals to a subthreshold defeat paradigm (total defeat time: ∼ 5s) (**Fig. 6k-m**), which was not sufficient to induce social avoidance during post-defeat SI test (**Fig. 6n-p**). Light delivery during the brief RI test to OXT^ChR2^ animals but not OXT^GFP^ males caused a significant decrease in the time spent around and investigating the cupped aggressor during the post-defeat SI test (**Fig. 6n-s**) although social fear, measured as immobility time, did not change significantly in any group (**Fig. 6t**). Thus, enhancing activity of SOR^OXT^ cells is sufficient to facilitate social avoidance learning after a mildly negative social experience.

## Discussion

To survive in a complex social group, it is important to learn to stay away from superior competitors. Indeed, a short 10-min defeat is sufficient to induce multi-week target-specific social avoidance^5^. Our study provides new mechanistic insight into the neural process that supports this rapid behavior change (**Fig. 6u**). Prior to defeat, aVMHvl^OXTR^ cells respond minimally to aggressor cues and the animals do not avoid the aggressor. During defeat, pain, likely caused by biting from the aggressor, evokes strong activation of oxytocin neurons in the SOR and presumably releases OXT, which then binds to OXTR to depolarize aVMHvl^OXTR^ cells and facilitates the potentiation of synapses that carry aggressor information. After defeat, when the animal encounters the aggressor again, due to the strengthened input that carries aggressor cues, aVMHvl^OXTR^ cells are now strongly activated, which in turn drives social avoidance to ensure the animals stay away from potentially disadvantageous conflicts (**Fig. 6u**).

Although aVMHvl^OXTR^ cells are indispensable for social avoidance, their role in social fear, as measured by immobility, could be less essential. While optogenetic activation of aVMHvl^OXTR^ cells induced both social avoidance and fear, inactivating the cells, knocking out aVMHvl OXTR, or ablating SOR^OXT^ cells reduced social avoidance but not social fear. Interestingly, acutely antagonizing OXTR in the aVMHvl during defeat reduced both social avoidance and social fear during post-defeat SI test. These results suggest that aVMHvl could be a part of the parallel circuits that drive social fear. While aVMHvl or its OXTR activation is acutely altered, it could affect acquisition and expression of social fear while long-term impairment of aVMHvl OXTR signaling, i.e. knock out and ablation, causes little behavioral deficits possibly due to compensatory effect of other pathways. These results also suggest that social fear and social avoidance, are two separable aspects of aversive social reactions and can be independently controlled at the neural circuit level. It is worth noting that SOR^OXT^-aVMHvl^OXTR^ is likely not the only circuit that supports social avoidance learning. Indeed, after blocking SOR^OXT^ or aVMHvl^OXTR^, most test animals still showed a low level of avoidance towards the aggressor although to a much lesser extent in comparison to control animals. We speculate that oxytocin coordinates OXTR cells in multiple brain regions to support this drastic and rapid learning process^41–44^.

As previously proposed, our results support a critical role of OXT in social behavior plasticity^45^. The findings that SOR^OXT^ cells are exclusively activated by painful stimuli and serve as a private source for aVMHvl^OXTR^ cells during defeat raises the possibility that there are distinct OXT subsystems dedicated for promoting social learning during positive and negative social encounters. Indeed, in contrast to the SOR^OXT^ cell responses, PVN^OXT^ cells are activated by positive social experience, such as gentle social touch, non-antagonistic social interaction and maternal shepherding^46–49^. Future studies that elucidate such OXT subsystems will be essential for harvesting the therapeutic potential of oxytocin in disorders with social deficits^50^. Finally, our studies suggest that aVMHvl, among many other hypothalamic regions^51^, is not only a crucial site for driving social behaviors but also a site of plasticity that mediates social experience-dependent behavioral changes.

## Methods

### Mice

All procedures were approved by the NYULMC Institutional Animal Care and Use Committee (IACUC) in compliance with the National Institutes of Health (NIH) Guidelines for the Care and Use of Laboratory Animals. Mice were housed under a 12 h light-dark cycle (dark cycle; 10 a.m. to 10 p.m. or 6:30 p.m. to 6:30 a.m.), with food and water available *ad libitum*. Oxtr^Cre^ (Strain #:031303)^24^, OXT^Cre^ (Strain #:024234)^34^, and Vglut2^Cre^ (Strain #:0169963)^52^ knock-in mice were purchased from Jackson Laboratory. OXTR^flox^ mice were kindly gifted by Dr. W. Scott Young III (National Institute of Mental Health) (now available from JAX, Strain #008471)^28^. Ai6 (Strain #:007906)^53^ mice were from Jackson Laboratory and crossed with Oxtr^Cre^ and Vglut2^Cre^ mice. Stimulus animals in RI test were BALB/c male (>9 weeks), C57BL/6N male and female mice (>8 weeks) originally purchased from Charles River and then bred in house, and Swiss Webster male and female mice (>11 weeks) purchased from Taconic, Charles River, or bred in house. After surgery with fiber or cannula implantation, all test mice were single housed.

### Viruses

The following AAVs were used in this study, with injection titers as indicated. AAV2 CAG-Flex-GCaMP6f-WPRE-SV40 (1.8 x 10^12^ vg/ml, UPenn, #V5747S) for fiber photometry was purchased from UPenn vector core. For the functional manipulation, the following AAVs were used. Optogenetic activation: AAV2 Ef1a-DIO-hChR2(H134R)-EYFP (4.2 × 10^12^ vg/ml, UNC, #AV4378); Chemogenetic inactivation: AAV2 hSyn-DIO-hM4Di-mCherry (1.5 × 10^13^ vg/ml, Addgene, #44362-AAV2); Optogenetic inactivation: AAV1 hSyn-SIO-stGtACR2-FusionRed (1.9 × 10^13^ vg/ml, Addgene, #105677-AAV1); OXTR knock-out: AAV1 CMV-HleGFP-Cre (1.1 × 10^13^ vg/ml, Addgene, #105545-AAV1) or AAV2 hSyn-GFP (3.4 × 10^12^ vg/ml, UNC, #4876D); Ablation of oxytocin cells: AAV8 hSyn-DIO-DTR (9.1 × 10^13^ vg/ml, Boston Children’s Hospital), AAV2 hSyn-DIO-GFP (4.0 × 10^12^ vg/ml, UNC, #4530C), or AAV2 hSyn-DIO-mCherry (1.8 × 10^13^ vg/ml, Addgene, #50459-AAV2). For slice electrophysiology, the following AAVs were used: AAV2 Ef1a-DIO-hChR2(H134R)-EYFP (4.2 × 10^12^ vg/ml, UNC, #AV4378) and AAV9 hSyn-ChrimsonR-tdTomato (2.6 × 10^13^ vg/ml, Addgene, #59171-AAV9).

### Drugs

For chemogenetic inhibition, 1 mg/kg Agonist21^27^ (Tocris, #5548) in saline was administered intraperitoneally. To block OXTRs in the aVMHvl, 250 nL/side 100 µM L-368,899 hydrochloride (Tocris, #2641) in saline was injected through implanted cannula bilaterally. To ablate OXT cells in the SOR, 50 µg/kg diphtheria toxin in saline (Sigma-Aldrich, #D0564) was administered intraperitoneally on two consecutive days (7 days before the pre-defeat SI test). A mixture of ketamine (100 mg/kg) and xylazine (10 mg/kg) in saline was administered intraperitoneally before perfusion.

### Stereotaxic surgery

Mice (8-12 weeks old) were anesthetized with 1-1.5% isoflurane and placed in a stereotaxic apparatus (Kopf Instruments Model 1900). Viruses were delivered into the targeted brain regions through glass capillaries using nanoinjector (World Precision Instruments, Nanoliter 2010) at a speed of 20 nL/min. 100-120 nL of AAV was injected into each targeted brain region. Stereotaxic injection coordinates were based on the Paxinos and Franklin mouse brain atlas^54^. For fiber photometry and optogenetic manipulation, polished optical fiber (440- or 230-µm diameter, Thorlabs) was implanted 150 or 200 µm above the virus injection site either immediately after virus injection or 2-3 weeks later. During the same surgery as optic fiber implantation, a 3D-printed head-fixation ring^55^ was cemented on the skull (C&B Metabond dental cement, Parkell) to allow head-fixation during fiber attach and detach, drug injection through cannula and head-fixed fiber photometry recording. Mice were single housed after optic fiber implantation. Histology was obtained from all test animals and only animals with correct virus expression and optic fiber placement were included in the final analysis.

### Common for all behavior tests

Mouse behaviors in all experiments were recorded from both the side and top of the cage using two synchronized cameras (Basler, asA640-100 gm and 120 gm) in a semi-dark room with infrared illumination. Video acquisition was achieved using StreamPix 5 (Noprix) at 25 frame/s. Manual behavioral annotation were performed on a frame-by-frame base using custom software written in MATLAB (https://pdollar.github.io/toolbox/)^19^. Most videos were annotated by people who did not perform behavior assay blindly and then checked and modified by an experimenter who was not blind to the animal’s group assignment. A subset of videos were also only annotated blindly. There is high consistency (around >90%) with annotations between performed blindly and not. During annotation, the neural responses were unknown to the experimenter. Custom DeepLabCut^56^-based models were constructed to track animals’ body center, head center and nose point in top-view videos. Movement velocity was calculated as the distance of an animal’s body center between two adjacent frames.

### Resident-intruder test

For inter-male RI tests with a goal to defeat the test animal, we introduced the test animal into the home cage of a sexually experienced, aggressive, and single-housed SW male mouse or introduced the SW male aggressor into the home cage of the test mouse (see individual experiments for details). For females, we introduced the test female into the home cage of a lactating female of SW or mixed background. The RI test typically lasted for 10 min although for several wild type male mice in Extended Data Fig. 1, the RI test was terminated after 5 min as the aggressor was highly aggressive and attacked the test animal continuously. Across experiments, each test mouse was defeated for approximately 20 s during the RI test. “Defeat” was annotated when the aggressor’s front end contacts the test animal’s back, presumably to bite, as the aggressor attacks the test mouse. “Fight” was annotated if the test animal successfully pushes or bites the aggressor when being attacked. For RI tests with non-aggressors, we introduced a Balb/C (BC) non-aggressive group-housed male mouse into the home cage of a male test mouse, and a C57 naïve female mouse into the home cage of a test female mouse. “Attack” was annotated when the test mouse initiates a suite of fast actions towards the non-aggressive intruder, including lunges, bites and tumbles. To analyze the behaviors, in addition to manual annotation, we also tracked the animals and calculated the percentage of time the test animal spent immobile (body center velocity <1.5 pixel/frame) during each minute of RI tests.

### Social interaction test

For SI test, the stimulus animals were always the same as those used in RI tests. In the aVMHvl^OXTR^ optogenetic activation experiment, group-housed non-aggressive Balb/C males were also used as stimulus animals. For females, on the day before the first SI test, the test animal was habituated to the empty metal wire cup (diameter of the cup bottom: 7.5 cm; height: 10.5 cm) in a clean cage for 10-15 min. There was no habituation for SI test in male. During the SI test, the stimulus animal was placed under a cup at one end of a clean cage with bedding. The test animal was then introduced into the cage and freely interacted with the cupped animal for 5 or 10 min. For each experiment, SI test was performed one day before and one day after RI test. For each experiment, the same set of aggressors were used for test and control groups to reduce the variability in defeat.

To analyze behaviors during SI test, we calculated the distance of the test animal’s body center to the cup center for each frame and constructed the histograms showing the distribution of test animal-cupped aggressor distance. The percentage of time an animal spent around the cup was calculated as the percentage of frames when the test animal to cup distance is <250 pixels, which is approximately half of the cage length. The investigation behavior was annotated manually as the time period when the test animal’s nose point is in close proximity to the cup. When the test animal stayed far from the cupped aggressor (distance >300 pixels) and when its body center movement velocity was <1.5 pixel/frame, the animal was considered as “immobile”. We then calculated the change index (CI) for each behavior parameter (P), including time around aggressor%, investigation time%, immobile time% as (P_after_ – P_before_)/ (P_after_ + P_before_). P_before_: value of the parameter before defeat; P_after_: value of the parameter after defeat. CI ranges from −1 to 1. Positive CI values indicate increases after defeat while negative CIs indicate decreases after defeat. Values close to −1 or 1 indicate large changes.

### Multi-animal social interaction test

For MSI test, the test arena (L × W × H: 22 inch × 18 inch × 16 inch) contained four wire meshed cup, one in each corner. On the habituation days (2 days), test animals freely explored the arena for around 20 min without any cups. On the test day, one cup was left empty and other three cups were each with a stimulus animal. The test animals were allowed to explore the arena freely for 10 min. MSI test was performed one the day before and one day after RI tests. The aggressor used in the MSI test were similar to those used in the SI tests. For each male MSI test, we also introduced a group-housed C57 male (16-24 weeks) and a Balb/C (BC) non-aggressive group-housed male (14-24 weeks), each under a cup, as stimulus animals. The same BC was encountered during the RI test during which the test animal either attacked or investigated it, but was never defeated by it. The C57 stimulus males were unfamiliar and only encountered during the MSI tests.

To analyze the behaviors, we calculated the distance between the animal’s head center to the center of each cup, when the distance is < 3× radius of the cup (*r_cup_*), the test animal was considered to be around the cupped animal. We also calculated the distance between animal’s nose point to the center of each cup, when the distance is <1.5× *r_cup_*, the test animal was considered as investigating the cupped animal. We then calculated the percentage of time each test animal spent on investigating, around the cup during before- and after-defeat MSI tests. We also calculated the average movement velocity when the test animal surrounds each cup.

### Optogenetic activation

To activate aVMHvl^OXTR^ cells, we injected 120 nL Cre-dependent ChR2 (control: GFP) expressing virus unilaterally into the aVMHvl (Bregma coordinates: AP, −1.455 mm; ML, −0.65 mm; DV, −5.70 mm) of OXTR^Cre^ mice and placed a 230-µm multi-mode optic fiber (Thorlabs, FT200EMT) 200 µm above injection site. On the test day, the implanted fiber was connected to a matching patch cord using a plastic sleeve (Thorlabs, ADAL1) to allow light delivery (Shanghai Dream Lasers). During the test, a non-aggressive BC male mouse and then an aggressive SW male mouse was placed under in a metal wire cup located at one end of a clean cage for around 15 min with 5 min in between. As for some test animals, assays with BC and SW male cup were performed on the different day with the same sequence of the stimulant animal (BC-SW). We then delivered 20 ms, 20 Hz, 1.5-2 mW light for 60 sec, followed by 0 mW sham light for 60s, and then repeated the light and sham trials once.

To activate SOR^OXT^ cells, we injected 120 nL Cre-dependent ChR2 (control: GFP) expressing virus bilaterally into the SOR (Bregma coordinates: AP, −1.36 mm; ML, ±0.90 mm; DV, −5.69 mm) of OXT^Cre^ male mice and placed two 230-µm multimode optic fibers (Thorlabs, FT200EMT) 200 µm above the injection sites. The behavior test started three weeks after virus injection. Each ChR2 and GFP animals went through two rounds of 3-days SI-RI-SI tests with at least 3 weeks in between. No light was delivered during SI tests. During RI test, a SW male aggressor was introduced into the home cage of the test mice until the aggressor defeated the test moue for 3-4 times with a total defeat duration of approximately 5 s. For one round of SI-RI-SI test, the light (3.5-4 mW, 20 ms, 20 Hz) was delivered during the entire RI test. For the other round of SI-RI-SI test, the test animals received no light during the RI tests. The order of light delivery during RI tests were counter-balanced across animals.

### Optogenetic and chemogenetic inhibition

To inactivate aVMHvl^OXTR^ cells, we injected 120 nL Cre-dependent stGtACR2 (control: mCherry) virus bilaterally into the aVMHvl (Bregma coordinates: AP, −1.455 mm; ML, ±0.65 mm; DV, −5.70 mm) of OXTR^Cre^ mice and placed two 230-µm multimode optic fibers (Thorlabs, FT200EMT) 200 µm above the injection sites. Three weeks after virus injection, all test animals went through 3-day SI-RI-SI tests. The SI tests lasted for 5 min instead of 10 min to reduce continuous light delivery duration. For each test animal, the aggressor was the same SW male mouse throughout the tests. No light was delivered during the first SI test or RI test. During post-defeat SIT test, one group of stGtACR2 animals received light (473nm, 3.5-4 mW, 20 ms, 20 Hz) for 5 min and the other group of stGtACR2 mice received no light. All mCherry animals received the light.

To chemogenetically inhibit aVMHvl^OXTR^ cells, we injected 100-110 nL Cre-dependent hM4Di (control: mCherry) virus bilaterally into the aVMHvl (Bregma coordinates: AP, −1.455 mm; ML, ±0.65 mm; DV, −5.70 mm) of OXTR^Cre^ mice. Three weeks later, animals went through 3-days SI-RI-SI tests with SW male aggressors. No drug was injected during the pre-defeat SI or RI tests. One hour before the post-defeat SI test, test animals were injected with 250 uL of saline or agonist 21 solution (1 mg/kg, Tocris, #5548) intraperitoneally.

### Knock-out of aVMHvl OXTRs

To knock out OXTR in the aVMHvl, we bilaterally injected 100 nL AAV expressing GFP-Cre (control: GFP) into the aVMHvl (Bregma coordinates: AP, −1.46 mm; ML, ±0.65 mm; DV, −5.70 mm) of OXTR^flox/flox^ male mice. 3-4 weeks after virus injection, all test animals went through the 3-day SI-RI-SI tests. During RI tests, SW aggressors were introduced into the home cage of the test animals for 10 minutes.

### OXTR antagonist application

To block aVMHvl OXTR, we implanted bilateral cannula (PlasticsOne, center-to-center distance: 1.5 mm) 0.7 mm above the aVMHvl (Bregma coordinates: AP, −1.45 mm; ML, ±0.75 mm; DV, −5.00 mm) of wild type C57BL/6 male mice. One week after the surgery, all animals went through the 3-day SI-RI-SI tests. To block aVMHvl OXTR during defeat, we injected 250 nL/side 100 µM L-368,899 hydrochloride (Tocris, #2641) into the aVMHvl through the cannula using a syringe (Hamilton, #65457-02) 20 min before RI test when the animal was head-fixed on a running wheel. To block OXTR during the post-defeat SI test, the same drug was injected 20 min before the SI test. Control animals were injected with saline and went through the same behavior tests. During the waiting time after drug injection, the animal was returned to its home cage. Before sacrificing the animals, we injected 250 nL 10 ng/mL DiI (ThermoFisher) to mark the site of injection.

### Ablation of OXT cells in the SOR

To ablate SOR^OXT^ cells, we injected 120 nL AAV expressing Cre-dependent DTR (control: GFP) into the SOR (Bregma coordinates: AP, −1.36 mm; ML, ±0.90 mm; DV, −5.69 mm) bilaterally using OXT^Cre^ male mice. Three weeks later, we intraperitoneally injected each animal with 50 µg/kg diphtheria toxin (DT, Sigma-Aldrich, #D0564) per day for two consecutive days. One week after the 1^st^ DT injection, all animals went through the 3-day SI-RI-SI tests.

### Fiber Photometry recording

We injected 120 nL AAV expressing Cre-dependent GCaMP6f into the SOR or aVMHvl unilaterally of 10-12-weeks old OXT^Cre^ and OXTR^Cre^ male and female mice. The following Bregma coordinates were used. Male SOR: AP, −1.36 mm; ML, −0.90 mm; DV, −5.69 mm from the top of the skull; female SOR: AP, −1.355 mm; ML, −0.88 mm; DV, −5.68 mm from skull surface. Male aVMHvl: AP, −1.46 mm; ML, −0.65 mm; DV, −5.7 mm from skull surface. Female aVMHvl: AP, −1.455 mm, ML, −0.645 mm; DV, −5.72 mm from skull surface. Recording started at least 3 weeks after virus injection.

Prior to fiber photometry recording, a ferrule sleeve (ADAL1-5, Thorlabs) was used to connect a matching patch cord to the implanted optic fiber when the animal was head fixed. For recordings, a 390-Hz sinusoidal 470-nm blue LED light (35 mW; LED light (M470F1, Thorlabs) driven by a LED driver (LEDD1B, Thorlabs) was bandpass-filtered (passing band: 472 ± 15 nm, Semrock, FF02-472/30-25) and delivered to the brain in to excite GCaMP6f. The emission light then passed through the same optic fiber, a bandpass filter (passing band: 534 ± 25 nm, Semrock, FF01-535/50), detected by a Femtowatt Silicon Photoreceiver (Newport, #2151) and recorded using RZ5 real-time processor (Tucker-Davis Technologies). The envelop of 390-Hz signals from the photoreceiver were extracted in real time using a custom-written program (Tucker-Davis Technologies) as the readout of GCaMP6f intensity. Top- and side-view behavior videos were simultaneously recorded (Basler, asA640-100 gm and 120 gm) and acquired using StreamPix 5 (Noprix) at 25 frame/s. Timestamps of video frames were used to align GCaMP6f signal and behaviors videos. For head-fixed recording of SOR^OXT^ cells, aggressor urine was collected from SW male aggressors or lactating SW female mice on the same day of recording. Urine was pooled from multiple aggressors including the mouse that defeated the test animal during the RI test. We then added 100 µL of urine to a Q-tip using a pipette and presented it to the recording animal manually for 10 sec with 50 sec in between. Male urine was presented to male test mice and female urine was presented to female animals. Gentle touch was performed onto the back of test animals with a large fluffy cotton ball (5 swipes for neck to tail base for each trial). Urine exposure and gentle touch were performed for 6 times for each animal. Back and tail pinches were applied with a pair of fine tweezers (FST, #91100-12) at a force that does not cause visible skin damage. Back pokes were applied using the pointy end of the same tweezers. 12 pinches or pokes were applied, each lasted for approximately 3 s, with 50-60 s in between.

To analyze the recording data, MATLAB function ‘‘msbackadj’’ with a moving window of 20% of the total recording duration was first applied to obtain the instantaneous baseline signal. The instantaneous Δ F/F was calculated as (F_raw_-F_baseline_)/F_baseline_. The Z scored ΔF/F of the entire recording session was calculated as (Δ F/F – mean(Δ F/F))/std(Δ F/F). The peri-event histogram (PETH) of Z scored Δ F/F aligned to a given behavior was constructed for each animal and then averaged across animals. The response during a specific behavior for each animal was calculated by averaging the Z scored Δ F/F during all time periods when the behavior occurred.

### Slice electrophysiology

To prepare brain slices for patch clamp recording, mice were anesthetized with isoflurane and brains were quickly removed and then immersed in ice-cold cutting solution for 1-2 min (in mM: 110 choline chloride, 25 NaHCO3, 2.5 KCl, 7 MgCl2, 0.5 CaCl2, 1.25 NaH2PO4, 25 glucose, 11.6 ascorbic acid and 3.1 pyruvic acid). The 275 μm aVMHvl coronal sections were cut using a Leica VT1200s vibratome, collected in oxygenated (95% O2 and 5% CO2) and pre heated (32–34 °C) artificial cerebrospinal fluid (ACSF) solution (in mM: 125 NaCl, 2.5 KCl, 1.25 NaH2PO4, 25 NaHCO3, 1 MgCl2, 2 CaCl2 and 11 glucose) and incubated for 30 min. The sections were then transferred to room temperature and continuously oxygenated until use.

Current and voltage whole cell patch clamp recordings were performed with micropipettes filled with intracellular solution containing (in mM: 145 K-gluconate, 2 MgCl2, 2 Na2ATP, 10 HEPES, 0.2 EGTA (286 mOsm, pH 7.2) or 135 CsMeSO3, 10 HEPES, 1 EGTA, 3.3 QX-314 (chloride salt), 4 Mg-ATP, 0.3 Na-GTP and 8 sodium phosphocreatine (pH 7.3 adjusted with CsOH)). Signals were recorded with MultiClamp 700B amplifier (Molecular Devices) and Clampex 11.0 software (Axon Instruments), digitized at 20 kHz with Digidata 1550B (Axon Instruments). After recording, data were analyzed using Clampfit (Molecular Devices) or MATLAB (Mathworks).

To characterize the physiological and synaptic properties of aVMHvl^OXTR^ cells, we identified ZsGreen positive cells in the aVMHvl on slices from OXTR^ZsGreen^ mice using an Olympus 40 × water-immersion objective with a GFP filter. For investigating intrinsic excitability, cells were recorded in current-clamp mode, and the number of action potentials was counted over 500-ms current steps. The current steps were consisted of 30 sweeps from −20 pA to 270 pA at 10 pA per step. sEPSCs and sIPSCs were recorded in the voltage-clamp mode. The membrane voltage was held at −70 mV for oEPSC recordings and at 0 mV for oIPSC recordings.

To investigate the efficacy of OXTR knockout, 120 nL AAV1-Cre-GFP and 120 nL AAV2-GFP were injected into the left (KO) and right (Control) sides of the brain, respectively. 3-4 weeks later, GFP positive cells in the aVMHvl from both KO and control sides were recorded in current clamp mode. All cells were recorded for 3-5 min after break-in until RMP was stable and then perfused with TGOT (250 µM) for 10 min. Cells that increased RMP for >4mV after 10 min TGOT perfusion were considered as OXTR positive.

In PVN^OXT^ and SOR^OXT^ optogenetic activation experiment, we injected 120-140 nL AAV2-Ef1a-DIO-ChR2-EYFP into either PVN or SOR of OXT^Cre^ male mice. After 4 weeks of virus incubation, aVMHvl cells were recorded in current-clamp mode. After the cell membrane potential was stabilized, we delivered 1 ms, 20 Hz blue light pulses (pE-300white; CoolLED) for 5 min to activate PVN^OXT^ or SOR^OXT^ processes. If the cell did not show a significant increase in RMP (>4mV) after light delivery, we then perfused TGOT (250 nM) in bath for 10 min to functionally determine the OXTR expression. The cells were separated into three categories based on their response to the light activation and TGOT perfusion: Light+, Light-TGOT+ and Light-TOGT-. Light+ and Light-TGOT+ cells were considered as putative OXTR expressing cells. Light-TGOT-cells were classified as OXTR-.

To examine the effect of OXT on synaptic transmission, we injected 100 nL AAV9-hSyn-ChrimsonR-tdTomato into PA of OXTR^ZsGreen^ male mice. After 4-week virus incubation, we obtained brain slices and performed current clamp recording of aVMHvl^OXTR^ cells. aVMHvl^OXTR^ cells are determined based on their ZsGreen expression. Before recording, the expression of ChrimsonR-tdTomato in PA was examined using an Olympus 40× water-immersion objective with a TXRED filter. The recording only proceeded if ChrimsonR-tdTomato was correctly and robustly expressed in the PA. To probe the excitatory synaptic responses of recorded aVMHvl^OXTR^ cells, we injected a small positive or negative current to keep the cell membrane potential around −70 mV and delivered a 1 ms 605-nm full-field light pulse every 10 s (0.1Hz) (pE-300white; CoolLED). After 5 min probing, we then delivered 3 trains of 20 Hz, 5 ms light pulses with 25 s per train and 5 s in between. During light pulse train delivery, the cells were not injected with any positive or negative current. After light train delivery, we again injected negative or positive current to maintain the membrane potential around −70 mV and probed the light-evoked EPSPs for 5 or 10 min, and then bath perfused 250 nM TGOT. After 10 min of TGOT perfusion, we delivered a second set of light pulse trains and then probed the light-evoked EPSPs for 5 or 10 min.

### Immunohistochemistry

For c-Fos and OXT staining, animals were deeply anesthetized with a mixture of ketamine (100 mg/kg) and xylazine (10 mg/kg) and transcardially perfused with 10 ml of PBS, followed by 10 ml of 4% paraformaldehyde in PBS. After perfusion, brains were harvested, soaked in 30% of sucrose in PBS for 24 hours at 4 ^ο^C and then embedded with O.C.T compound (Fisher Healthcare). 40 μm thick coronal brain sections were cut using a cryostat (Leica). Brain sections were washed with PBST (0.3% Triton X-100 in PBS, 10 min), blocked in 5% normal donkey serum (NDS, Jackson Immuno Research) in PBST for 30 min at room temperature (RT), and then incubated with primary antibodies in 5% NDS in PBST overnight at RT (about 18 hours). Sections were then washed with PBST (3×10 min), incubated with secondary antibodies in 5% NDS in PBST for 4 hours at RT, washed with PBST (2×10 min) and DAPI-mixed (1:10000, Thermo Scientific) PBS solution (1×20 min). Slides were coverslipped using mounting medium (Fluoromount, Diagnostic BioSystems) after drying.

The primary antibodies used were: rabbit anti-Oxytocin (1:5000, Immunostar, #20068, Lot #1607001), guinea pig anti-c-Fos (1:2000, Synaptic Systems, 226-005, Lot #2-10, 2-13), rabbit anti-Vasopressin (1:5000, Immunostar, #20069, Lot #1004001), and anti-GFP (1:2000, abcam, ab13970, lot#GR3190550-2). The secondary antibodies used were: Cy3-AffiniPure donkey anti-rabbit IgG (1:500, Jackson Immuno Research, 711-165-152, lot#124528), Cy5-AffiniPure donkey anti-rabbit IgG (1:250, Jackson Immuno Research, 711-175-152, lot#150312), Alexa Fluor 488-conjugated goat anti-guinea pig IgG (1:500, Invitrogen. #A11073, lot#2160428), or Alexa Fluor 488-conjugated donkey anti-chicken IgY(IgG) (1:500, Jackson Immuno Research. 703-545-155, lot#116967). The 10x or 20× fluorescent images were acquired to determine the overall expression pattern in each brain regions by Olympus VS120 Automated Slide Scanner and its specific software OlyVIA. The 20× fluorescent confocal images were acquired by Zeiss LSM 800 and its specific software (Zeiss, ZEN 2.3 system) for cell counting.

### *In situ* hybridization

To prepare the sections for *in situ* hybridization (ISH), 10-12-weeks-old C57BL/6 male mice were anesthetized with a mixture of ketamine (100 mg/kg) and xylazine (10 mg/kg) and transcardially perfused with 10 ml of DEPC treated PBS (DEPC-PBS), followed by 10 ml of 4% paraformaldehyde in DEPC-PBS (PFA, from paraformaldehyde 32% solution, Electron Microscopy Sciences). After perfusion, brains were harvested, soaked in 30% of sucrose in DEPC-PBS for 24 hours at 4 ^ο^C and then embedded with O.C.T compound (Fisher Healthcare). 30 μm thick coronal brain sections were cut using a cryostat (model #CM3050S, Leica). The sections were placed on MAS-coated glass slides (MAS-03, Matsunami) and stored at −80 ^ο^C before use.

To synthesize the cDNA for the *OXT* and *OXTR* probes, their original templates were from mouse brain cDNA (cDNA-mmu-01, BiOSETTIA). cDNA was amplified by PCR methods using the following oligo-DNA primers and the products were purified with micro spin columns (MACHEREY-NAGEL, 74060910). Each reverse primer also possesses T3 sequence for transcription.

OXT1-forward: TGGCTTACTGGCTCTGACCT

OXT1-reverse: AATTAACCCTCACTAAAGGGAGGAAGCGCGCTAAAGGTAT

OXTR1-forward: GGCGGTCCTGTGTCTCATAC

OXTR1-reverse: AATTAACCCTCACTAAAGGGCTCCACATCTGCACGAAGAA

OXTR2-forward: TTCATCATTGCCATGCTCTT

OXTR2-reverse: AATTAACCCTCACTAAAGGGGGGTGGCTCTCATTTCCTTT

OXTR3-forward: GCTGGAGATAGGAGGCAGTG

OXTR3-reverse: AATTAACCCTCACTAAAGGGGCTGTGTCACTCACCAGACG

The *OXT* probe are approximately 400 bp in length and the *OXTR* probes are approximately from 750 bp to 1000 bp in length. ISH probes were prepared by *in vitro* transcription with DIG RNA Labeling Mix (Roche Applied Science, #11277073910) or Fluorescein RNA Labeling mix (Roche Applied Science, #11685619910) and T3 polymerase (Roche Applied Science, #11031163001). *OXT* probe was labeled with Flu and *OXTR* probes were labeled with DIG.

Brain sections including the VMHvl underwent ISH at 56 °C overnight. After a series of post-hybridization washing and blocking, Flu-positive cells were visualized with anti-FITC antibody (PerkinElmer, #NEF710001EA, 1:200 in blocking buffer) followed by TSA biotin amplification reagent (PerkinElmer, #NEF749A001KT, 1:100 in 1 × plus amplification diluent) and streptavidin Alexa488 (Invitrogen, #S11223, 1:250 in blocking buffer). DIG-positive cells were visualized with anti-DIG antibody (Roche Applied Science, #11207733910, 1:250 in blocking buffer) and TSA Cy3 amplification regent (PerkinElmer, #NEL744001KT, 1:100 in 1 × plus amplification diluent). Sections were counterstained with 4’,6-diamino-2-phenylindole dihydrochloride (DAPI, 1:10000 in PBS, Thermo Scientific) and mounted with a cover glass using Fluoromount (Diagnostic BioSystems, #K024).

The 20× fluorescent images were acquired using a slide scanner (Olympus VS120). 20× fluorescent confocal images were acquired using Zeiss LSM 800 (Zeiss, ZEN 2.3 system).

### Quantification and statistical analysis

Statistical analyses were performed using MATLAB (version 2019b or 2021b, Mathworks) and Prism9 (GraphPad Software, RRID: SCR_002798). All statistical analyses were two-tailed. Parametric tests, including paired t-test, unpaired t-test and one-way ANOVA were used if distributions passed Kolmogorov–Smirnov or Shapiro-Wilk tests for normality or else nonparametric tests, including Wilcoxon matched-pairs signed rank test, Mann-Whitney test, Kruskal-Wallis test and Friedman test were used. For comparisons across multiple groups and variables, two-way ANOVA were used without formally testing the normality of data distribution. Post-hoc multi-pair comparison tests were performed after one-way ANOVA, Kruskal-Wallis test, Friedman test and two-way ANOVA. *p< 0.05; **p<0.01; ***p<0.001; ****p<0.0001. Error bars represent ± SEM. For detailed statistical results, see Figure legends.

### Data and code availability

Data reported in this paper and code used in this paper are available from the authors upon request.

## Acknowledgements

We thank Robert C. Froemke and György Buzsáki for helpful discussion and comments and Jing-Jing Liu for assisting annotation of behavior assays and Ryo Kenmochi for cartoon illustrations and all the members in U19 Oxytocin and Dayu Lin lab for valuable discussion and comments. This research was supported by NIH grants U19NS107616 (R. W. T., A. C. M., and D. L.), R01MH71739 (R. W. T.), R01MH101377, 1R01HD092596, and U01NS113358 (D. L.); the Mathers Foundation (D. L.); the Vulunerable Brain Project (D. L.); and the Uehara Memorial Foundation, JSPS Overseas Research Fellowship, and Osamu Hayaishi Memorial Scholarship (T. O.).

## Author contributions

T. O. and D. L. conceived the project. D. L. supervised the study, designed experiments, and performed data analysis. T. O. designed and performed most of behavioral experiments, and immunostaining, and analyzed the data. R. Y. performed all slice recording experiments and analyzed the data. Y. J. performed most of the behavior experiments in female mice and some fiber photometry recordings. D. W. performed preliminary slice recording experiments. B. D. performed automated behavioral analysis. R. T., G. Z., and C.-X. W. assisted with behavior annotation. X. W. evaluated OXTR-KO efficiency under the supervision of R. W. T.. A. C. M. performed preliminary behavioral analysis. T. O. and D. L. wrote the paper with inputs from other authors.

---

**Extended Data Fig. 1:**
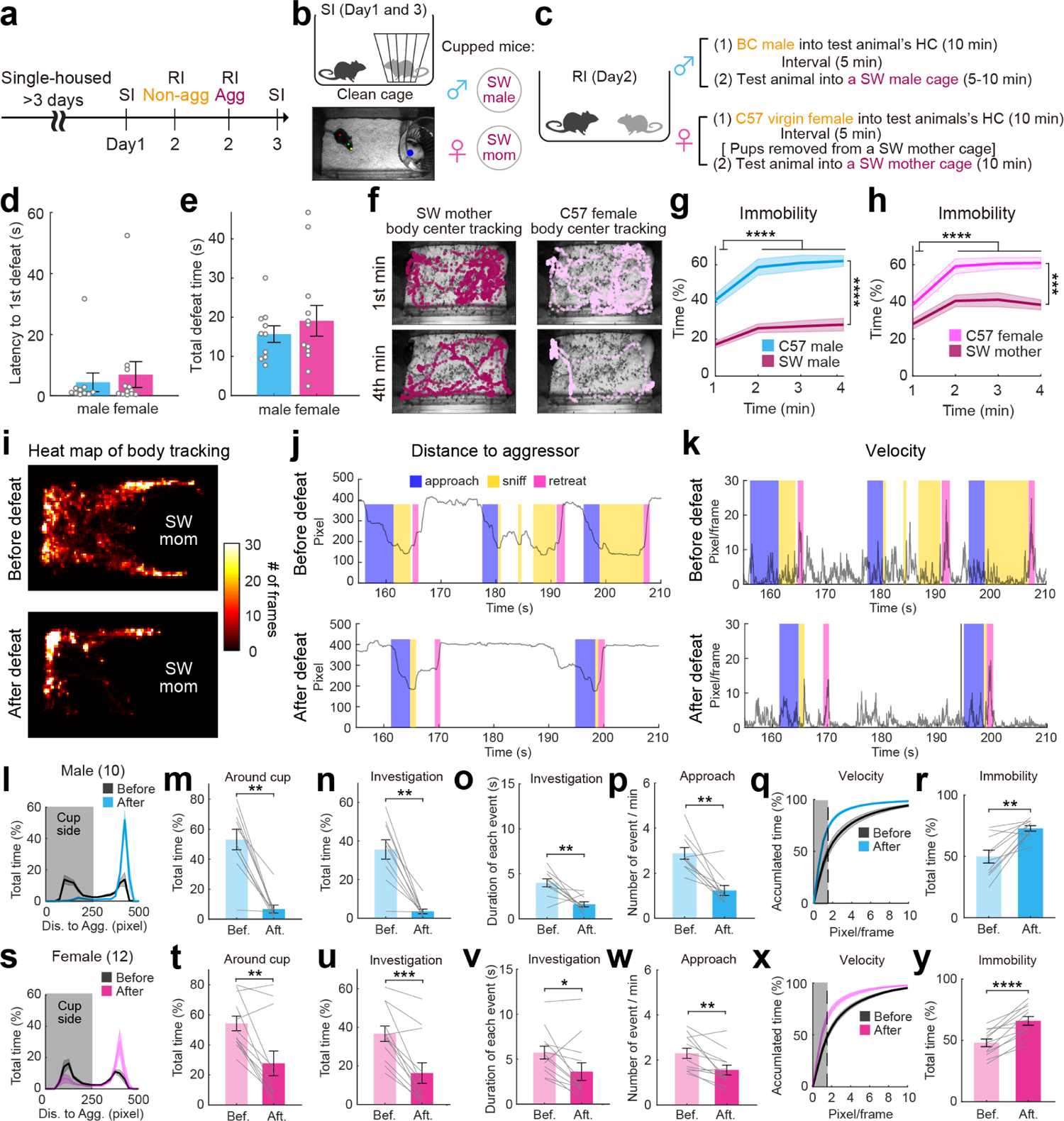
10 min social defeat is sufficient to induce social avoidance in male and female mice. **a,** Experimental timeline. **b,** Cartoon illustration of the social interaction (SI) test (top) and a video frame overlaid with DLC tracking results (bottom). Red dot: body center; green dot: head center; yellow dot: nose point; blue dot: cup center. **c**. Cartoon illustration and timeline of the resident-intruder (RI) test. **d,** The latency to first defeat in male and female mice. **e,** The total defeat duration during the 10-min RI test in male and female mice. **f,** Video frames from SI tests, overlaid with the movement trajectories of the SW mother aggressor (maroon) and C57 test female (pink) during the 1^st^ (top) and 4^th^ (bottom) minute of the test. **g-h,** The percentage of time the male (**g**) and female (**h**) test mice and the aggressors showing immobility over the course of RI tests. **i,** Heatmaps showing the body center location of a representative female mouse during pre-defeat and post-defeat SI tests. **j-k,** Representative traces showing the distance between the test animal body center to the cup center (**j**) and the movement velocity of the test animal (**k**) in pre- and post-defeat SI tests. Color shades indicate manually annotated behaviors. **l and s,** Distribution of the distance between the test animal’s body center and cup center during the pre-defeat (gray) and post-defeat (color) SI tests for males (**l**) and females (**s**). Shades shown in gray represent the distance range considered as “around cup”. **m and t,** The percentage of total time the male (**m**) and female (**t**) test mice spent around the aggressor cup (distance < 250 pixels) during SI tests. **n and u,** The percentage of total time the male (**n**) and female (**u**) test mice spent on investigating the aggressor cup during SI tests. **o and v,** The average duration of each investigation episode of the male (**o**) and female (**v**) test mice during SI tests. **p and w,** The cup approach frequency of the male (**p**) and female (**w**) test mice during SI tests. **q and x,** Accumulative plots showing the distribution of movement velocity when the male (**q**) and female (**x**) test mice are far away (distance >300 pixels) from the cupped aggressor during the pre-defeat (gray) and post-defeat (color) SI tests. Gray shades mark immobility zones (pixel/frame<1.5). **r and y,** The percentage of time the male (**r**) and female (**y**) test mice spent immobile when far from the cupped aggressor during SI tests. Shades in **g, h, l, q, s,** and **x**, and error bars in **d, e, m-p, r, t-w,** and **y** represent ± SEM. Circles and lines in **d, e, m-p, r, t-w** and **y** represent individual animals. n=10 male mice in **d, e, g,** and **l-r**. n =12 female mice in **d, e** and **s-y** and 9 females in **h.** Statistical analyses were performed with Mann Whitney test (**d**), unpaired t-test (**e**), two-way repeated measure ANOVA with Sidak’s multiple comparisons test (**g, h**), Wilcoxon matched-pairs signed rank test (**m, n, p, t, u, v**), and paired t-test (**o, r, w, y**). All statistical analyses were two-tailed. *p<0.05, **p<0.01, ***p<0.001, and ****p<0.0001.

**Extended Data Fig. 2:**
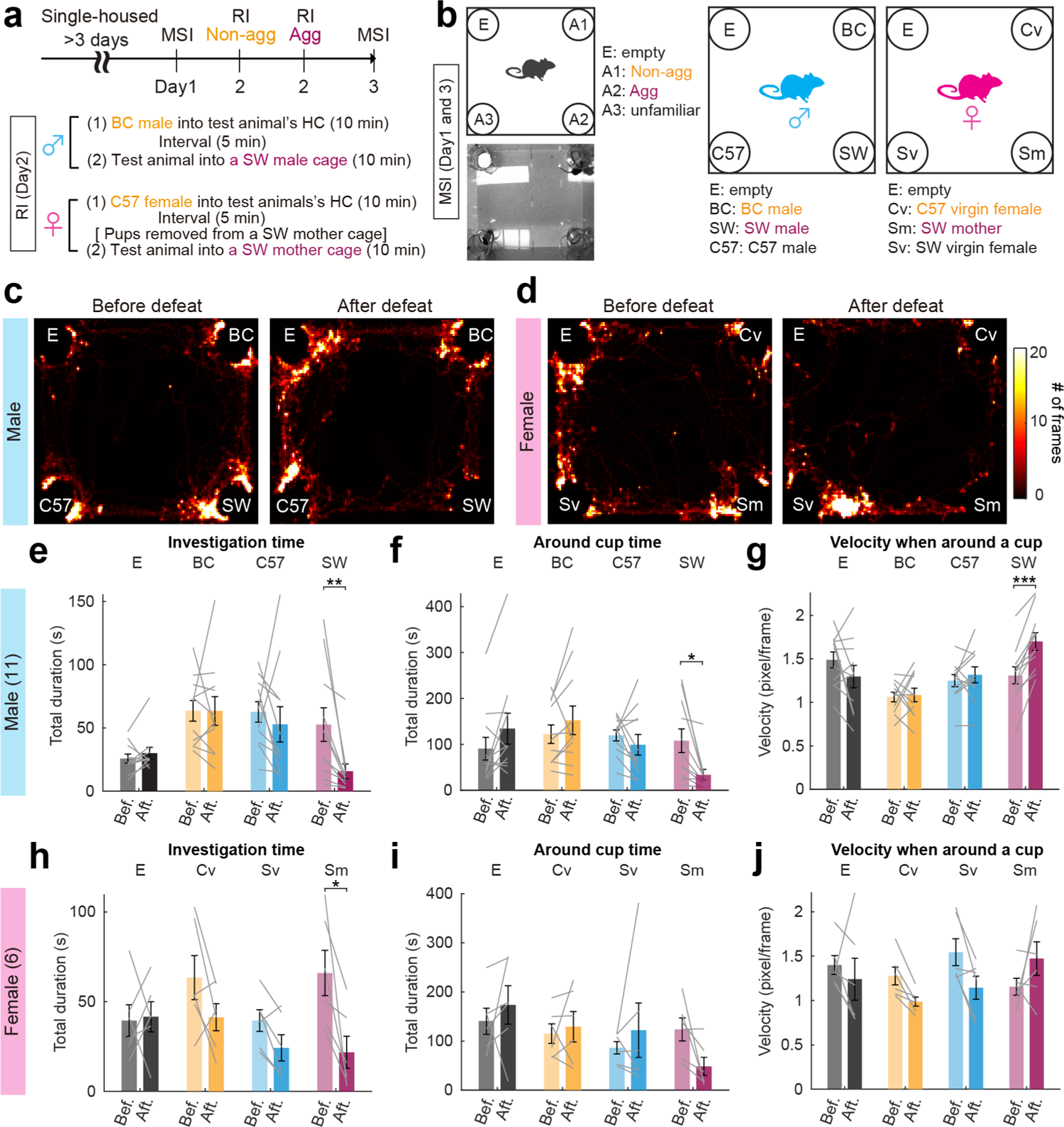
Defeated animals show avoidance specifically towards the winner of the fight. **a,** Experimental timeline. **b,** Cartoon illustration, a snapshot of the multi-animal social interaction (MSI) test, and stimulus animals used for male and female test animals. **c-d,** Heatmaps showing the body center location of a representative male (**c**) and a female (**d**) mouse in MSI tests before and after defeat. **e-f,** Total time male test mice spent on investigating (**e**) and around (**f**) each cupped animal during MSI tests before and after defeat. **g**, Movement velocity of the test male mice when around each cup during MSI tests before and after defeat. **h-j,** Data from female mice. Plots follow the convention of **e-g**. Error bars in **e-j** represent ± SEM. Lines in **e-j** represent individual animals. Two-way repeated measure ANOVA with Sidak’s multiple comparisons test was performed. All statistical analyses were two-tailed. *p<0.05, **p<0.01, and ***p<0.001.

**Extended Data Fig. 3:**
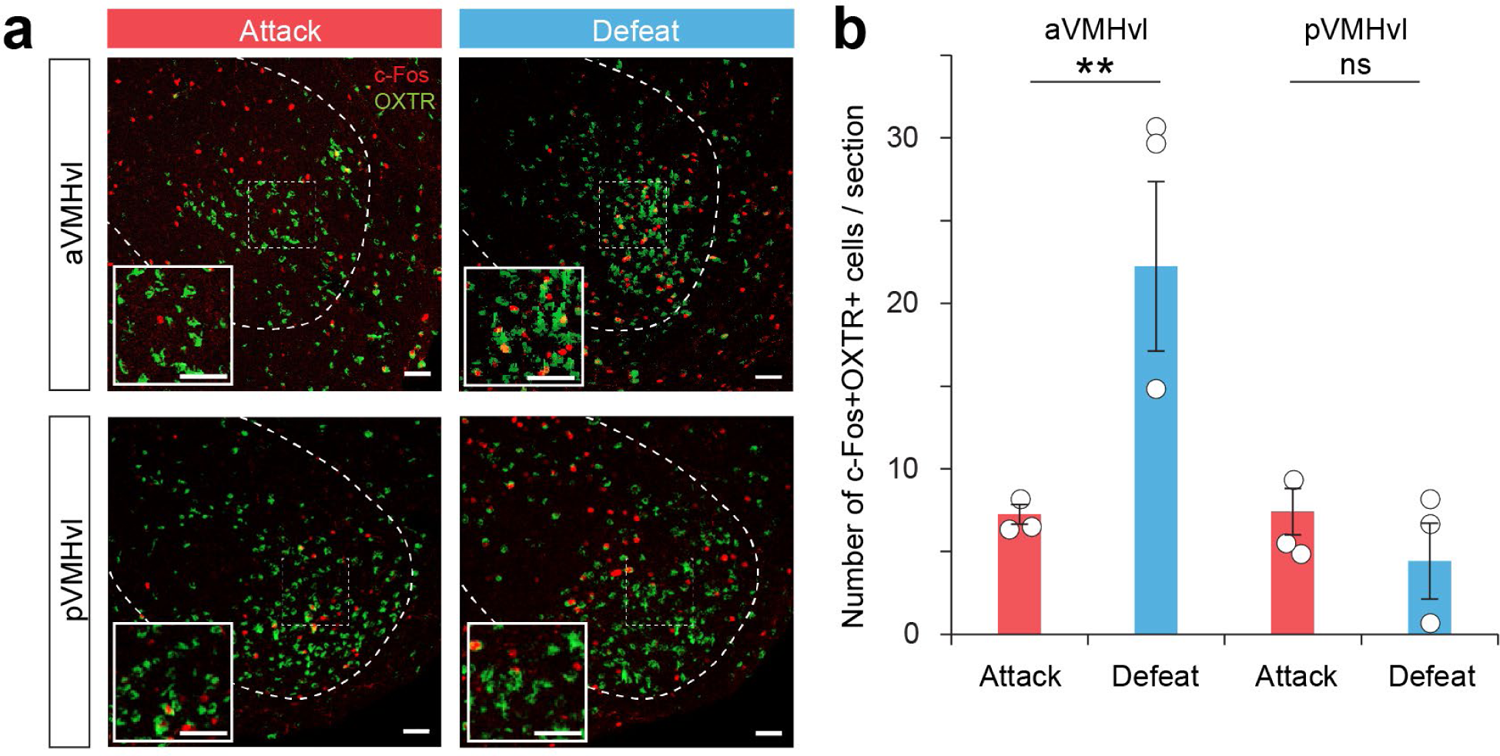
Defeat induced high c-Fos expression in aVMHvl^OXTR^ cells. **a,** Representative histological images showing c-Fos (red) and OXTR (OXTR^ZsGreen^, green) expression in the aVMHvl (Bregma: −1.48 mm) and pVMHvl (Bregma: −1.84 mm) in OXTR^Cre^ × Ai6 (OXTR^ZsGreen^) male mice after attack or social defeat. Insets showing enlarged views of the boxed areas. Dashed lines mark the boundary of the VMH. Scale bars: 50 µm. **b,** The number of c-Fos and OXTR double positive cells after attack and defeat in the aVMHvl and pVMHvl. Error bars: ± SEM. Circles represent individual animals. Two-way repeated measure ANOVA with Sidak’s multiple comparisons test. **p<0.01. ns: not significant.

**Extended Data Fig. 4:**
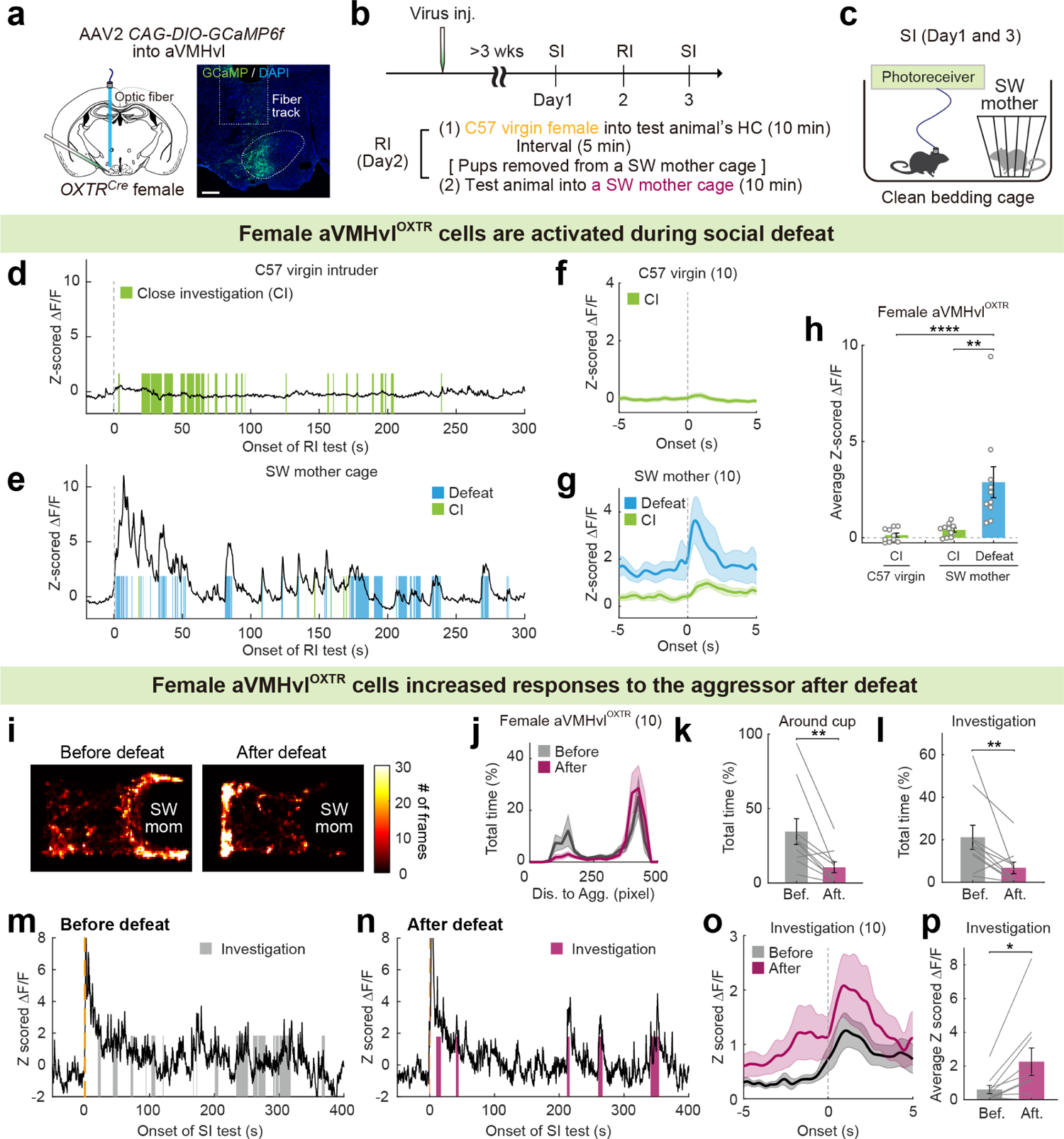
Female aVMHvl^OXTR^ cells increase activity to the aggressor after defeat. **a,** Schematics of virus injection and a representative histology image. Dashed lines mark the boundary of the aVMH. Scale bar: 200 µm. **b,** Experimental timeline. **c,** Cartoon illustration of SI test. **d-e,** Representative Z scored GCaMP6f traces from a recording female mouse during RI tests with a non-aggressive naïve C57 (**d**) and an aggressive lactating SW (**e**) female mouse. **f-g,** PETHs of Z scored GCaMP6f signals aligned to onset of investigating C57 intruders (**f**), and investigating and being defeated by SW residents (**g**). **h,** Average Z scored ΔF/F of aVMHvl^OXTR^ cells during various social behaviors in RI tests. **i,** Heatmaps showing the body center location of a representative test female during pre-defeat and post-defeat SI tests. **j,** Distribution of the distance between the test animal’s body center and cup center during the pre- and post-defeat SI tests. **k,** The percentage of total time the test mice spent around the aggressor cup during pre- and post-defeat SI tests. **l,** The percentage of total time the test mice spent on investigating the aggressor cup during pre- and post-defeat SI tests. **m-n,** Representative Z scored GCaMP6f traces from a recording mouse during pre-defeat (**m**) and post-defeat (**n**) SI tests. Shades represent investigation events. **o,** PETHs of Z scored GCaMP6f signals aligned to onset of investigating the cupped aggressor during pre- and post-defeat SI tests. **p,** Average Z scored ΔF/F of aVMHvl^OXTR^ cells during investigating aggressor in the pre- and post-defeat SI tests. Shades in **f, g, j** and **o,** and error bars in **h, k, l** and **p** represent ± SEM. Circles and lines in **h, k, l** and **p** represent individual animals. n=10 female mice in **f-h, j-l** and **o-p**. Statistical analyses were performed with Kruskal-Wallis test with Dunn’s multiple comparisons test (**h**), Wilcoxon matched-pairs signed rank test (**k, l**) and paired t-test (**p**). All statistical analyses were two-tailed. *p<0.05, **p<0.01, and ****p<0.0001.

**Extended Data Fig. 5:**
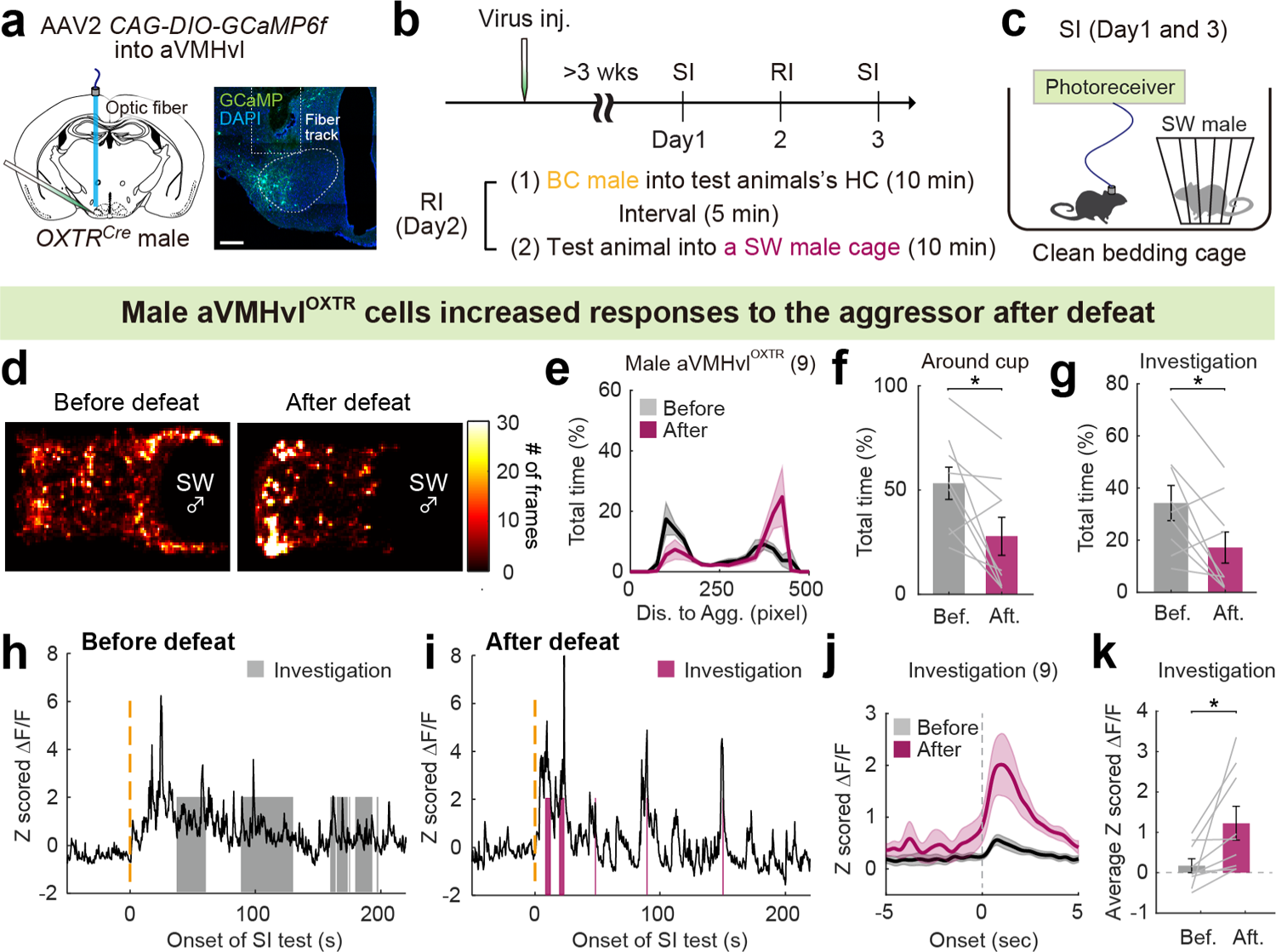
aVMHvl^OXTR^ cells increase activity to the aggressor after defeat in male mice. **a,** Schematics of virus injection a representative histology image. Dashed line marks the aVMH. Scale bar: 200 µm. **b,** Experimental timeline. **c.** Cartoon illustration of SI test. **d,** Heatmaps showing the body center location of a representative test male during pre- and post-defeat SI tests. **e,** Distribution of the distance between the test animal’s body center and cup center during the pre- and post-defeat SI tests. **f,** The percentage of total time test mice spent around the aggressor cup during pre- and post-defeat SI tests. **g,** The percentage of total time the test mice spent on investigating the cupped aggressor during pre- and post-defeat SI tests. **h-i,** Representative Z scored GCaMP6f traces from a recording male mouse during pre-defeat (**h**) and post-defeat (**i**) SI tests. Shades represent investigation events. **j,** PETHs of Z scored GCaMP6f signals aligned to onset of investigating aggressor. **k,** Average Z scored ΔF/F of aVMHvl^OXTR^ cells during investigating aggressor in pre- and post-defeat SI tests. Shades in **e** and **j**, and error bars in **f, g** and **k** represent ± SEM. Lines in **f, g** and **k** represent individual animals. n=9 male mice in **e-g, j** and **k**. Statistical analyses were performed with Wilcoxon matched-pairs signed rank test (**g**) and two-tailed paired t-test (**f, k**). *p<0.05.

**Extended Data Fig. 6:**
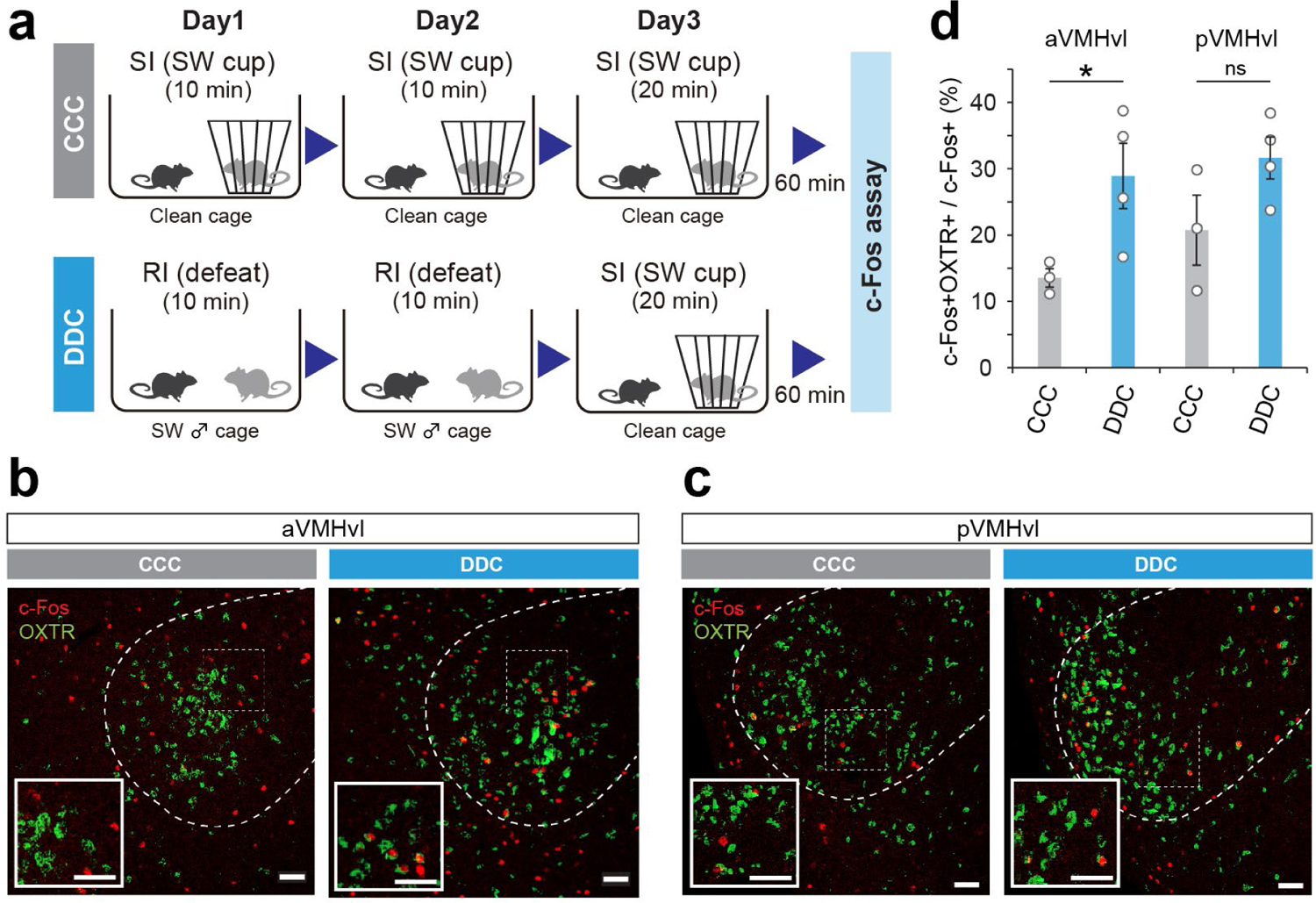
Defeat experience enhances aggressor cue-induced c-Fos in aVMHvl^OXTR^ cells during subsequent encounters. **a,** Experimental design. CCC: SI (Cup)-SI (Cup)-SI (Cup) (top); DDC: RI (Defeat)-RI(Defeat)-SI (Cup) (bottom). **b and c,** Representative images showing the expression of OXTR (OXTR^ZsGreen^, green) and c-Fos (red) in the aVMHvl (**b**) and pVMHvl (**c**) in animals experienced CCC or DDC. Insets showing enlarged views of the boxed areas in the aVMHvl and pVMHvl. Dashed lines mark the boundary of the VMH. Scale bars: 50 µm. **d,** The percentage of CCC and DDC induced c-Fos cells that express OXTR in the aVMHvl and pVMHvl. n=3 (CCC) and 4 (DDC) male mice. Error bars: ± SEM. Circles indicate individual animals. Two-way repeated measure ANOVA with Sidak’s multiple comparisons test. *p<0.05. ns: not significant.

**Extended Data Fig. 7:**
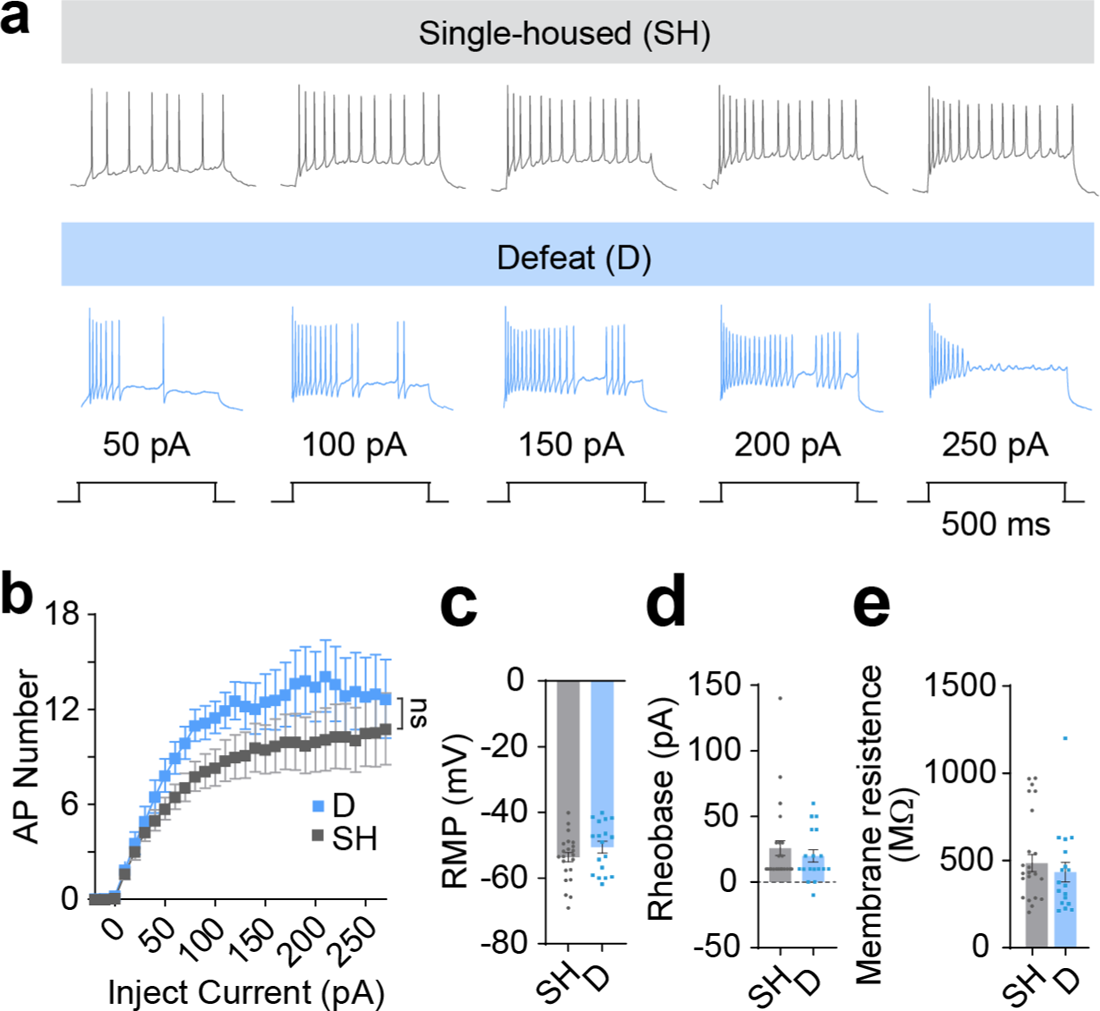
No change in excitability of aVMHvl^OXTR^ cells one day after defeat. **a,** Representative recording traces of aVMHvl^OXTR^ cells under specific current steps, ranging from 50 pA to 250 pA, from single-housed (SH) and defeated (D) male mice. **b,** Frequency-current (F-I) curve of aVMHvl^OXTR^ cells in SH and D groups. **c,** Resting Membrane Potential (RMP) of aVMHvl^OXTR^ cells in SH and D groups. **d,** Rheobase of aVMHvl^OXTR^ cells in SH and D groups. **e,** Membrane resistance of aVMHvl^OXTR^ cells in SH and D groups. Error bars in **b-e** represent ± SEM. Squares in **c-e** represent individual recording cells. n= 27 (SH) and 18 (D) cells, each from 3 male mice, in **b-e**. Statistical analyses were performed with two-way repeated measure ANOVA (**b**), and unpaired t-test (**c-e**).

**Extended Data Fig. 8:**
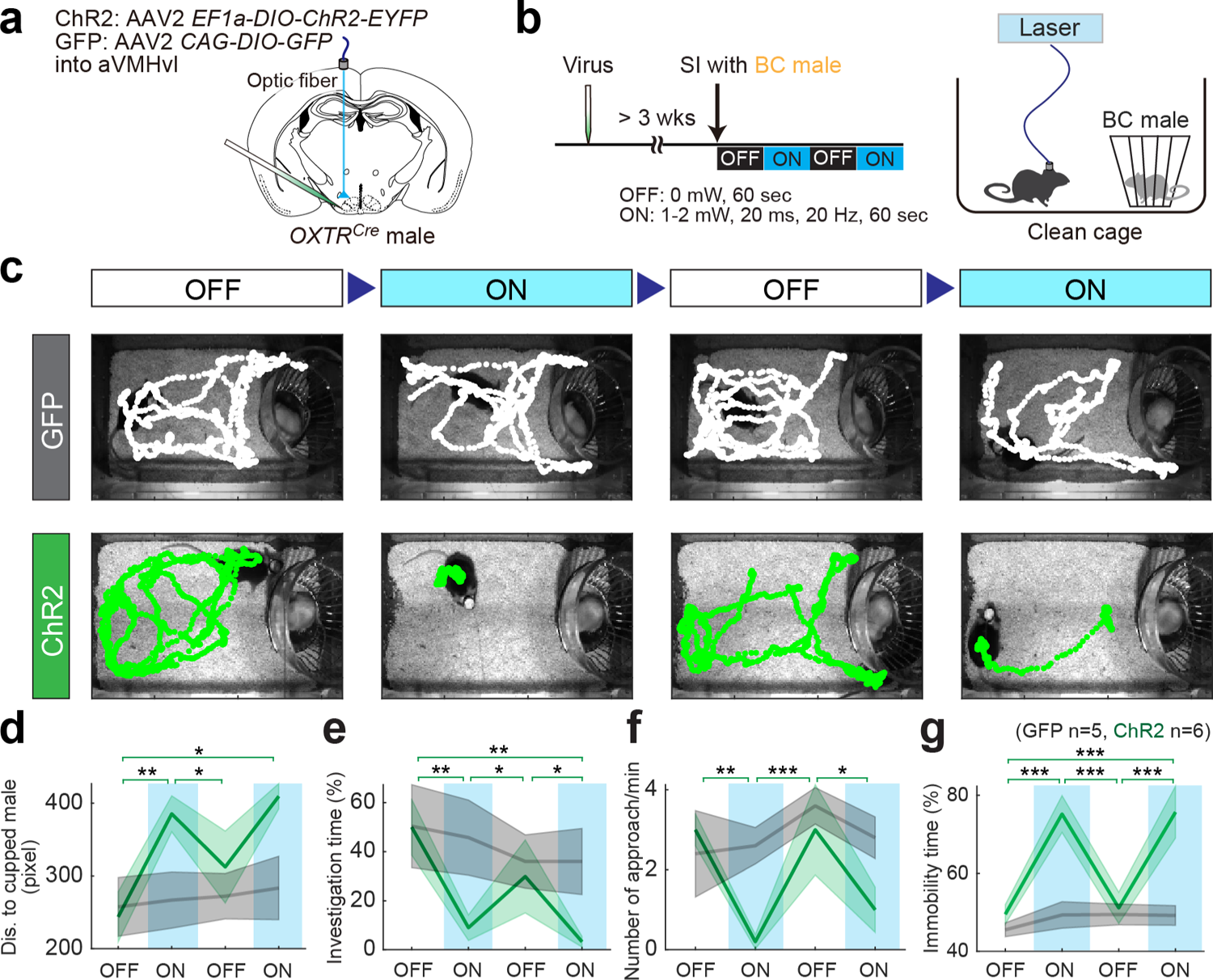
Optogenetic activation of aVMHvl^OXTR^ cells induces social avoidance and fear towards a non-aggressor. **a,** Virus injection schematics. **b,** Experimental timeline and light delivery protocol. **c,** Video frames from SI tests overlaid with movement trajectories of a GFP (gray) and a ChR2 animal (green) during interleaved light (ON) and a sham (OFF) stimulation trials. **d-g,** Average distance to the cupped male (**d**), percentage of time spent on investigating the cupped BC male (**e**), frequency of approaching the cup (**f**) and percentage time the test animal spent immobile (**g**) during light-on (blue) and light-off periods in GFP (gray) and ChR2 (green) groups. Statistical results were between ON and OFF periods in ChR2 animals. No significant difference between ON and OFF periods in GFP animals. n=5 (GFP) and 6 (ChR2) male mice. Shades in **d-g** represent ± SEM. Statistical analyses were performed with two-way RM ANOVA with Holm-Sidak’s multiple comparisons test (**d-g**). All statistical analyses were two-tailed. *p<0.05, **p<0.01, and ***p<0.001.

**Extended Data Fig. 9:**
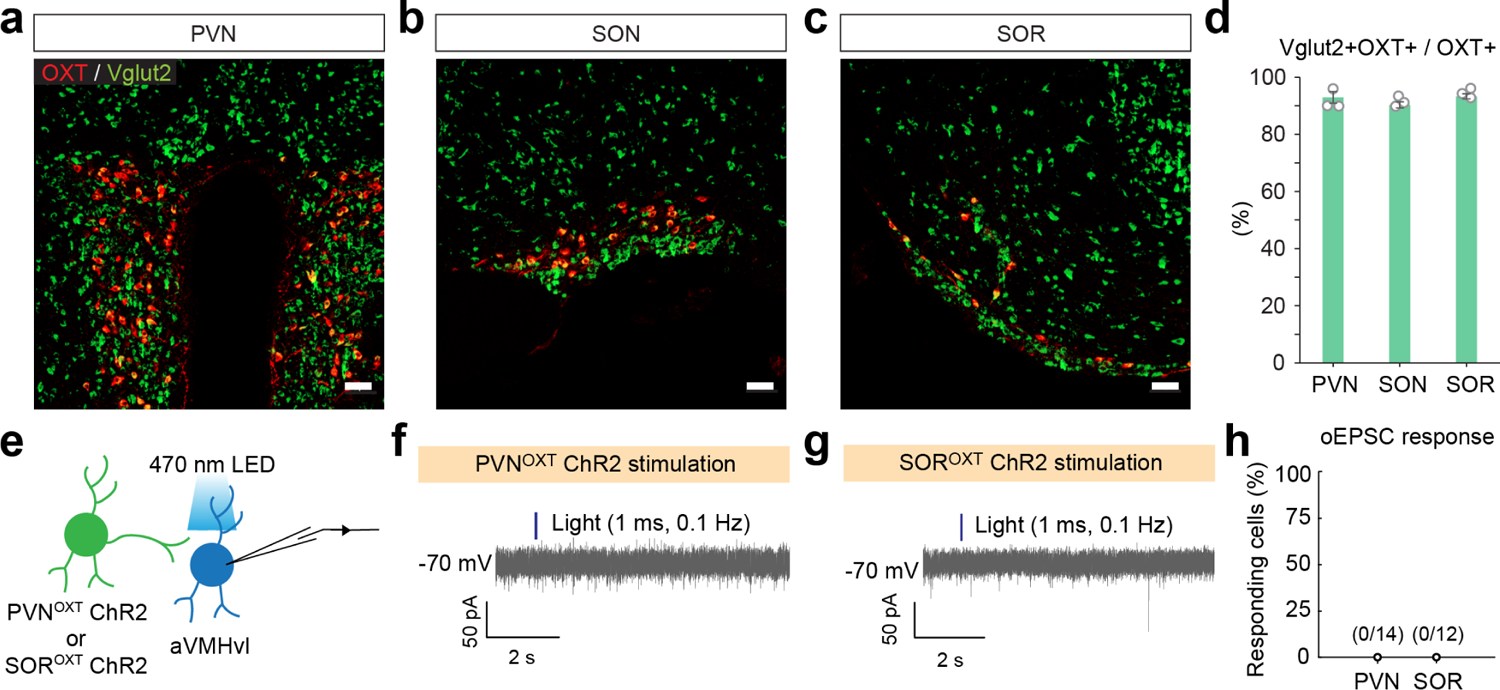
OXT cells are largely glutamatergic but do not provide glutamatergic input to aVMHvl^OXTR^ cells. **a-c,** Histology images showing oxytocin (OXT, red) immunostaining and Vglut2 (green) expression in the PVN (**a**), SON (**b**), and SOR (**c**) from Vglut2^Cre^ × Ai6 male mice. Scale bars: 50 µm. **d.** The percentage of OXT-positive cells that express Vglut2 in the PVN, SON and SOR. Circles represent individual animals. Error bars: ± SEM. n=3 male mice. **e.** Slice recording schematics. **f-g,** Representative voltage clamp recording traces from aVMHvl^OXTR^ cells when a 1 ms light pulse (blue vertical bar) was delivered to activated PVN^OXT^ (**f**) or SOR^OXT^ (**g**) input. **h.** None of the aVMHvl^OXTR^ cells showed light evoked EPSC during PVN^OXT^ (0/14 cells) or SOR^OXT^ optogenetic activation (0/12 cells).

**Extended Data Fig. 10:**
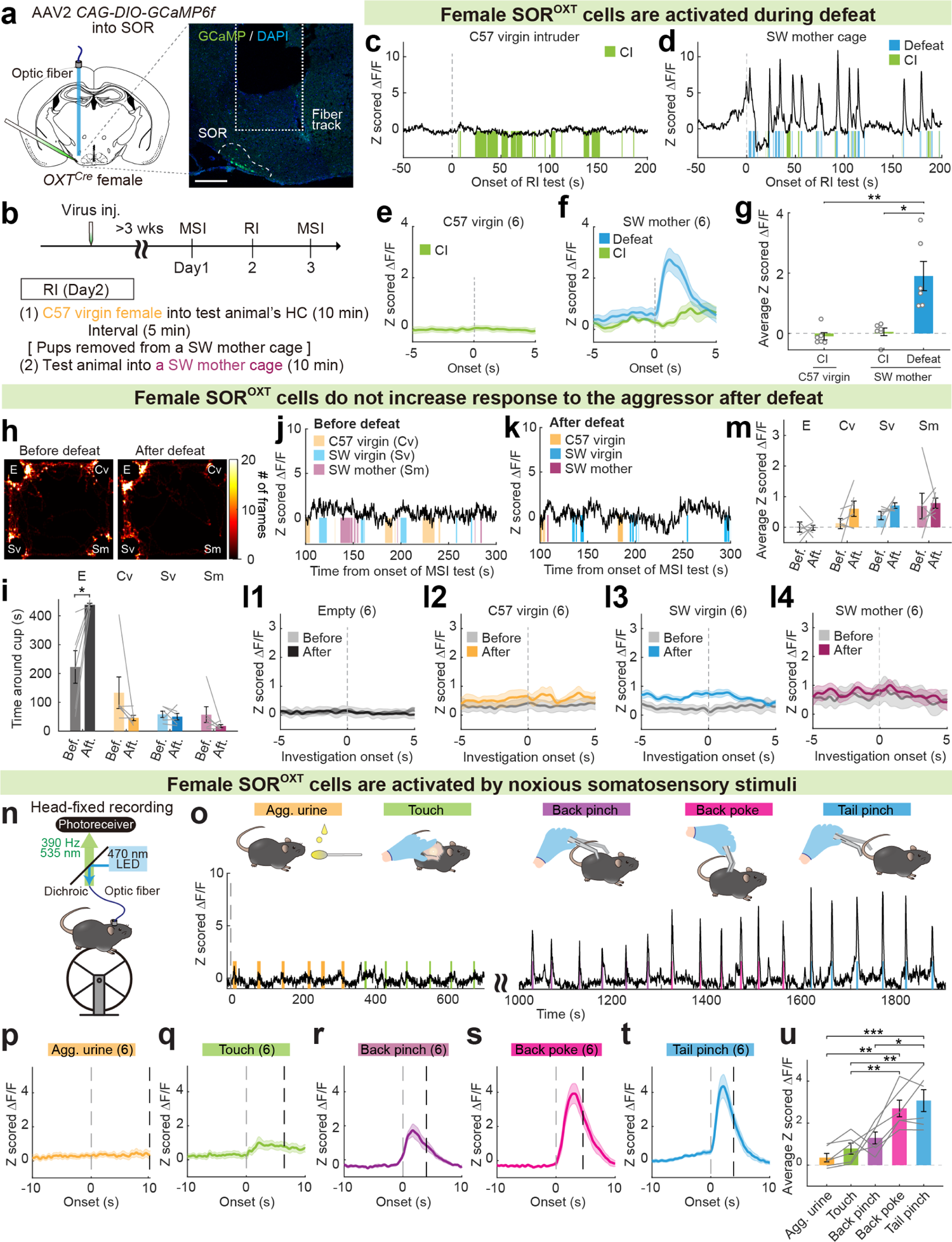
SOR^OXT^ cells in female mice are activated by noxious stimuli experienced during defeat. **a,** Schematics of virus injection and a representative histological image for fiber photometry recording of SOR^OXT^ cells in female mice. Dashed line marks SOR. Scale bar: 200 µm. **b,** Experimental timeline. **c-d,** Representative Z scored GCaMP6f traces of SOR^OXT^ cells from an animal that encountered a naïve C57BL/6 female intruder (**c**) or a SW lactating female mouse in the SW cage (**d**). **e-f**, PETHs of Z scored GCaMP6f signals aligned to close investigation (CI) of C57BL/6 female intruders (**e**), investigating and being defeated by SW mothers (**f**). **g,** Average of Z scored ΔF/F of SOR^OXT^ cells during various social behaviors. **h,** Heatmaps showing the body center location of a recording mouse in MSI tests before and after defeat. E: empty; Cv: familiar C57BL/6 virgin female; Sv: unfamiliar virgin SW female; and Sm: SW mother. **i,** Time spent around each cup during MSI tests before and after defeat. **j-k,** Representative Z scored GCaMP6f traces from a female recording mouse during pre-defeat and post-defeat MSI tests. Shades represent investigation periods of different cupped stimulus animals. Periods investigating the empty cup were not marked. **l,** PETHs of Z scored GCaMP6f signals aligned to onset of investigation of different cupped stimuli. Gray: pre-defeat; Color: post-defeat. **m,** Average of Z scored ΔF/F responses of SOR^OXT^ cells during investigation of various cupped stimuli in the pre-defeat and post-defeat MSI tests. **n,** Schematics of head-fixed fiber photometry recording of SOR^OXT^ cells. **o,** Representative Z scored GCaMP6f trace of SOR^OXT^ cells during delivery of aggressor urine on a Q-tip, gentle touch, back pinch, back poke and tail pinch. **p-t,** PETHs of Z scored GCaMP6f signals aligned to onset of aggressor urine presentation (**p**), gentle touch (**q**), back pinch (**r**), back poke (**s**), and tail pinch (**t**). **u,** Average Z scored ΔF/F during various stimulus delivery. Shades in **e-f, l** and **p-t**, and error bars in **g, i, m and u** represent ± SEM. Circles in **g** and lines in **i, m** and **u** represent individual animals. n = 6 female mice in **e-g, i, l, m, p-u**. Statistical analyses were performed with Kruskal-Wallis test with Dunn’s multiple comparisons test (**g**), two-way repeated measure ANOVA with Sidak’s multiple comparisons test (**i, m**), and one-way repeated measure ANOVA with Tukey’s multiple comparisons test (**u**), All statistical analyses were two-tailed. *p<0.05, **p<0.01, and ***p<0.001.

## References

1. Archer, J. The behavioural biology of aggression. (Cambridge University Press, 1988).

2. Crowcroft, P. Mice all over. (Chicago Zoological Society, 1973).

3. Blanchard, R. J., Blanchard, D. C., Takahashi, T. & Kelley, M. J. Attack and defensive behaviour in the albino rat. Animal Behaviour 25, 622–634 (1977).

4. Martinez, M., Calvo-Torrent, A. & Pico-Alfonso, M. A. Social defeat and subordination as models of social stress in laboratory rodents: a review. Aggressive Behavior 24, 241–256 (1998).

5. Qi, C. C. et al. Interaction of basolateral amygdala, ventral hippocampus and medial prefrontal cortex regulates the consolidation and extinction of social fear. Behav Brain Funct 14, 7, doi:10.1186/s12993-018-0139-6 (2018).

6. Silva, B. A. et al. Independent hypothalamic circuits for social and predator fear. Nature neuroscience 16, 1731–1733, doi:10.1038/nn.3573 (2013).

7. Faturi, C. B., Rangel, M. J., Jr., Baldo, M. V. & Canteras, N. S. Functional mapping of the circuits involved in the expression of contextual fear responses in socially defeated animals. Brain Struct Funct 219, 931–946, doi:10.1007/s00429-013-0544-4 (2014).

8. Schlund, M. W. et al. Human social defeat and approach-avoidance: Escalating social-evaluative threat and threat of aggression increases social avoidance. J Exp Anal Behav 115, 157–184, doi:10.1002/jeab.654 (2021).

9. Banks, R. (ERIC Clearinghouse on Elementary and Early Childhood Education, University …, 1997).

10. Huhman, K. L. et al. Conditioned defeat in male and female Syrian hamsters. Horm Behav 44, 293–299 (2003).

11. Markham, C. M., Taylor, S. L. & Huhman, K. L. Role of amygdala and hippocampus in the neural circuit subserving conditioned defeat in Syrian hamsters. Learn Mem 17, 109–116, doi:10.1101/lm.1633710 (2010).

12. Day, D. E., Cooper, M. A., Markham, C. M. & Huhman, K. L. NR2B subunit of the NMDA receptor in the basolateral amygdala is necessary for the acquisition of conditioned defeat in Syrian hamsters. Behav Brain Res 217, 55–59, doi:10.1016/j.bbr.2010.09.034 (2011).

13. Markham, C. M., Luckett, C. A. & Huhman, K. L. The medial prefrontal cortex is both necessary and sufficient for the acquisition of conditioned defeat. Neuropharmacology 62, 933–939 (2012).

14. Sakurai, K. et al. Capturing and Manipulating Activated Neuronal Ensembles with CANE Delineates a Hypothalamic Social-Fear Circuit. Neuron 92, 739–753, doi:10.1016/j.neuron.2016.10.015 (2016).

15. Wang, L. et al. Hypothalamic Control of Conspecific Self-Defense. Cell Rep 26, 1747–1758 e1745, doi:10.1016/j.celrep.2019.01.078 (2019).

16. Diaz, V. & Lin, D. Neural circuits for coping with social defeat. Curr Opin Neurobiol 60, 99–107, doi:10.1016/j.conb.2019.11.016 (2020).

17. Krzywkowski, P., Penna, B. & Gross, C. T. Dynamic encoding of social threat and spatial context in the hypothalamus. Elife 9, doi:10.7554/eLife.57148 (2020).

18. Newman, S. W. The medial extended amygdala in male reproductive behavior. A node in the mammalian social behavior network. Ann N Y Acad Sci 877, 242–257, doi:10.1111/j.1749-6632.1999.tb09271.x (1999).

19. Lin, D. et al. Functional identification of an aggression locus in the mouse hypothalamus. Nature 470, 221–226, doi:10.1038/nature09736 (2011).

20. Yang, C. F. et al. Sexually dimorphic neurons in the ventromedial hypothalamus govern mating in both sexes and aggression in males. Cell 153, 896–909, doi:10.1016/j.cell.2013.04.017 (2013).

21. Toth, I. & Neumann, I. D. Animal models of social avoidance and social fear. Cell Tissue Res 354, 107–118, doi:10.1007/s00441-013-1636-4 (2013).

22. Gandelman, R. Mice: postpartum aggression elicited by the presence of an intruder. Horm Behav 3, 23–28, doi:10.1016/0018-506x(72)90003-7 (1972).

23. Nasanbuyan, N. et al. Oxytocin-Oxytocin Receptor Systems Facilitate Social Defeat Posture in Male Mice. Endocrinology 159, 763–775, doi:10.1210/en.2017-00606 (2018).

24. Daigle, T. L. et al. A Suite of Transgenic Driver and Reporter Mouse Lines with Enhanced Brain-Cell-Type Targeting and Functionality. Cell 174, 465–480 e422, doi:10.1016/j.cell.2018.06.035 (2018).

25. Boyden, E. S., Zhang, F., Bamberg, E., Nagel, G. & Deisseroth, K. Millisecond-timescale, genetically targeted optical control of neural activity. Nature neuroscience 8, 1263–1268, doi:10.1038/nn1525 (2005).

26. Mahn, M. et al. High-efficiency optogenetic silencing with soma-targeted anion-conducting channelrhodopsins. Nat Commun 9, 4125, doi:10.1038/s41467-018-06511-8 (2018).

27. Thompson, K. J. et al. DREADD Agonist 21 Is an Effective Agonist for Muscarinic-Based DREADDs in Vitro and in Vivo. ACS Pharmacol Transl Sci 1, 61–72, doi:10.1021/acsptsci.8b00012 (2018).

28. Lee, H. J., Caldwell, H. K., Macbeth, A. H., Tolu, S. G. & Young, W. S., 3rd. A conditional knockout mouse line of the oxytocin receptor. Endocrinology 149, 3256–3263, doi:10.1210/en.2007-1710 (2008).

29. Liao, P. Y., Chiu, Y. M., Yu, J. H. & Chen, S. K. Mapping Central Projection of Oxytocin Neurons in Unmated Mice Using Cre and Alkaline Phosphatase Reporter. Front Neuroanat 14, 559402, doi:10.3389/fnana.2020.559402 (2020).

30. Rhodes, C. H., Morrell, J. I. & Pfaff, D. W. Immunohistochemical analysis of magnocellular elements in rat hypothalamus: distribution and numbers of cells containing neurophysin, oxytocin, and vasopressin. J Comp Neurol 198, 45–64, doi:10.1002/cne.901980106 (1981).

31. Castel, M. & Morris, J. F. The neurophysin-containing innervation of the forebrain of the mouse. Neuroscience 24, 937–966, doi:10.1016/0306-4522(88)90078-4 (1988).

32. Ludwig, M. Dendritic release of vasopressin and oxytocin. J Neuroendocrinol 10, 881–895, doi:10.1046/j.1365-2826.1998.00279.x (1998).

33. Pow, D. V. & Morris, J. F. Dendrites of hypothalamic magnocellular neurons release neurohypophysial peptides by exocytosis. Neuroscience 32, 435–439, doi:10.1016/0306-4522(89)90091-2 (1989).

34. Wu, Z. et al. An obligate role of oxytocin neurons in diet induced energy expenditure. PLoS One 7, e45167, doi:10.1371/journal.pone.0045167 (2012).

35. Kim, D.-W. Multimodal analysis of cell types in a hypothalamic node controlling social behavior in mice, California Institute of Technology, (2020).

36. Klapoetke, N. C. et al. Independent optical excitation of distinct neural populations. Nat Methods 11, 338–346, doi:10.1038/nmeth.2836 (2014).

37. Yamaguchi, T. et al. Posterior amygdala regulates sexual and aggressive behaviors in male mice. Nature neuroscience 23, 1111–1124, doi:10.1038/s41593-020-0675-x (2020).

38. Stagkourakis, S., Spigolon, G., Liu, G. & Anderson, D. J. Experience-dependent plasticity in an innate social behavior is mediated by hypothalamic LTP. Proceedings of the National Academy of Sciences of the United States of America 117, 25789–25799, doi:10.1073/pnas.2011782117 (2020).

39. Zha, X. et al. VMHvl-Projecting Vglut1+ Neurons in the Posterior Amygdala Gate Territorial Aggression. Cell Rep 31, 107517, doi:10.1016/j.celrep.2020.03.081 (2020).

40. Malinow, R. & Miller, J. P. Postsynaptic hyperpolarization during conditioning reversibly blocks induction of long-term potentiation. Nature 320, 529–530, doi:10.1038/320529a0 (1986).

41. Zoicas, I., Slattery, D. A. & Neumann, I. D. Brain oxytocin in social fear conditioning and its extinction: involvement of the lateral septum. Neuropsychopharmacology 39, 3027–3035, doi:10.1038/npp.2014.156 (2014).

42. Ebner, K., Wotjak, C. T., Landgraf, R. & Engelmann, M. A single social defeat experience selectively stimulates the release of oxytocin, but not vasopressin, within the septal brain area of male rats. Brain Res 872, 87–92, doi:10.1016/s0006-8993(00)02464-1 (2000).

43. Williams, A. V. et al. Social approach and social vigilance are differentially regulated by oxytocin receptors in the nucleus accumbens. Neuropsychopharmacology 45, 1423–1430, doi:10.1038/s41386-020-0657-4 (2020).

44. Menon, R. et al. Oxytocin Signaling in the Lateral Septum Prevents Social Fear during Lactation. Curr Biol 28, 1066–1078 e1066, doi:10.1016/j.cub.2018.02.044 (2018).

45. Froemke, R. C. & Young, L. J. Oxytocin, Neural Plasticity, and Social Behavior. Annu Rev Neurosci 44, 359–381, doi:10.1146/annurev-neuro-102320-102847 (2021).

46. Carcea, I. et al. Oxytocin neurons enable social transmission of maternal behaviour. Nature 596, 553–557, doi:10.1038/s41586-021-03814-7 (2021).

47. Yu, H. et al. Social touch-like tactile stimulation activates a tachykinin 1-oxytocin pathway to promote social interactions. Neuron 110, 1051–1067 e1057, doi:10.1016/j.neuron.2021.12.022 (2022).

48. Tang, Y. et al. Social touch promotes interfemale communication via activation of parvocellular oxytocin neurons. Nature neuroscience 23, 1125–1137, doi:10.1038/s41593-020-0674-y (2020).

49. Resendez, S. L. et al. Social Stimuli Induce Activation of Oxytocin Neurons Within the Paraventricular Nucleus of the Hypothalamus to Promote Social Behavior in Male Mice. J Neurosci 40, 2282–2295, doi:10.1523/JNEUROSCI.1515-18.2020 (2020).

50. Erdozain, A. M. & Penagarikano, O. Oxytocin as treatment for social cognition, not there yet. Frontiers in Psychiatry 10, 930 (2020).

51. Wei, D., Talwar, V. & Lin, D. Neural circuits of social behaviors: Innate yet flexible. Neuron 109, 1600–1620, doi:10.1016/j.neuron.2021.02.012 (2021).

52. Vong, L. et al. Leptin action on GABAergic neurons prevents obesity and reduces inhibitory tone to POMC neurons. Neuron 71, 142–154, doi:10.1016/j.neuron.2011.05.028 (2011).

53. Madisen, L. et al. A robust and high-throughput Cre reporting and characterization system for the whole mouse brain. Nature neuroscience 13, 133–140, doi:10.1038/nn.2467 (2010).

54. Franklin, K. B. J. & Paxinos, G. Paxinos and Franklin’s The mouse brain in stereotaxic coordinates. Fourth edition. edn, (Academic Press, an imprint of Elsevier, 2013).

55. Osborne, J. E. & Dudman, J. T. RIVETS: a mechanical system for in vivo and in vitro electrophysiology and imaging. PLoS One 9, e89007, doi:10.1371/journal.pone.0089007 (2014).

56. Mathis, A. et al. DeepLabCut: markerless pose estimation of user-defined body parts with deep learning. Nature neuroscience 21, 1281–1289, doi:10.1038/s41593-018-0209-y (2018).

